# Structural connectivity shapes a cross-modal gradient of brain flexibility

**DOI:** 10.64898/2025.12.29.696703

**Authors:** Matteo Neri, Marianna Angioielli, Elsa Bourgue, Camille Mazzara, Emahnuel Troisi Lopez, Matteo Demuru, Mario De Luca, Enrica Gallo, Mario Quarantelli, Andrea Brovelli, Pierpaolo Sorrentino

## Abstract

The brain must rapidly and dynamically recruit the appropriate brain regions to respond to environmental changes and sustain flexible cognition. An extensive body of literature views the ability to spontaneously explore a rich repertoire of activity patterns, even during rest, as a correlate of the capacity to respond rapidly to internal fluctuations and external stimuli. Moreover, brain flexibility at rest successfully indexes pathological conditions such as Parkinson’s disease, multiple sclerosis, and amyotrophic lateral sclerosis. However, previous work provides no information about the contribution of each region to the flexibility of brain dynamics. Given the functional and structural heterogeneity of brain regions, we hypothesized their roles in sustaining flexibility to be heterogeneous and related to regional structural properties. To systematically investigate these hypotheses, we define the flexibility gradient (FG), a new measure, which quantifies regional contributions to flexibility. We employed three datasets spanning different recording techniques and a range of spatial and temporal scales: 47 MEG recordings, 11 EEG recordings, and 77 fMRI recordings along with the corresponding tractographies. We found that FG is not homogeneously distributed, with associative and occipital regions contributing the most. This gradient is symmetric across left and right hemispheres, and the structural properties of brain regions correlate with their contribution to flexibility. Our findings are stable across modalities and temporal scales, highlighting that brain flexibility is not only a global dynamical property but also a spatially structured feature shaped by the anatomical organization of the brain.

## INTRODUCTION

To sustain a broad range of cognitive functions, the brain must dynamically orchestrate its activity, exploring a vast repertoire of states and efficiently recruiting the functional systems required for a given task. A growing body of literature has shown that this capacity is reflected in the richness of activation patterns observed in functional brain data during rest^1–5^. More specifically, both the size and the variability of this repertoire have been linked to the brain’s ability to flexibly and appropriately engage its functional resources to sustain flexible cognition^6–8^.

To investigate the flexibility of large-scale brain dynamics, a new approach has recently been proposed that focuses on the analysis of transient bursts of brain activity^1^. This approach is supported by a growing body of evidence showing that novel insights into brain dynamics can be gained by studying nonlinear, intermittent bursts that propagate across the brain along spatiotemporal trajectories^6,9,10^. Along these lines, the flexibility of the dynamics has been described as the number of (unique) spatial configurations with which these bursts spread across the brain. It is posited that a healthy brain would maximise the number of visited spatial configurations (i.e., activity patterns)^2^, possibly by dwelling in a near-critical regime. In this line of work, a shrinkage of the functional repertoire has been described in several neurological diseases, proportional to, or directly predictive of, clinical disabilities^1,2^. Furthermore, recent work has shown that changes across physiological states, such as the phases of the menstrual cycle, are also mirrored by changes in flexibility^3^.

However, the role of individual brain regions in attaining flexible large-scale dynamics, while likely highly heterogeneous, remains largely unknown. For instance, what is the contribution of a given brain region to the flexibility of large-scale brain dynamics? How does the structural connectivity of a region relate to its contribution to flexibility? These questions are particularly relevant in light of evidence for disease-specific spatial patterns and the pronounced structural and functional heterogeneity observed across brain regions. Moreover, a growing body of literature highlights the role of structural connectivity in shaping time-averaged functional interactions^10–15^. In light of these findings, we hypothesize that the contributions of individual brain regions to dynamic flexibility are highly heterogeneous across the cortex, and that a region’s centrality within the structural connectivity network plays a key role in determining its ability to contribute to flexibility.

To address these questions, we analyzed brain activities across multiple temporal scales and spatial resolutions using three complementary datasets: source-reconstructed MEG from 47 healthy adults, EEG from 11 healthy adults, and a publicly available resting-state fMRI dataset from 77 healthy subjects (Figure 1a). We systematically investigated how individual brain regions contribute to the flexibility of large-scale neural dynamics, based on the hypothesis that the structural heterogeneity of brain regions shapes their contributions^10,13,15^. To this end, we introduce the **Flexibility Gradient (FG)**, which measures the reduction in the brain’s functional repertoire when a given region is removed. We hypothesised that FG would organise across brain lobes in a hemispherically symmetric, hierarchical topography that reflects the connectome’s structural architecture. To probe this hypothesis, we used subject-specific structural connectivity matrices. We related the FG and the nodal degree across brain regions, using data collected with different spatial and temporal resolutions.

**Figure 1.**
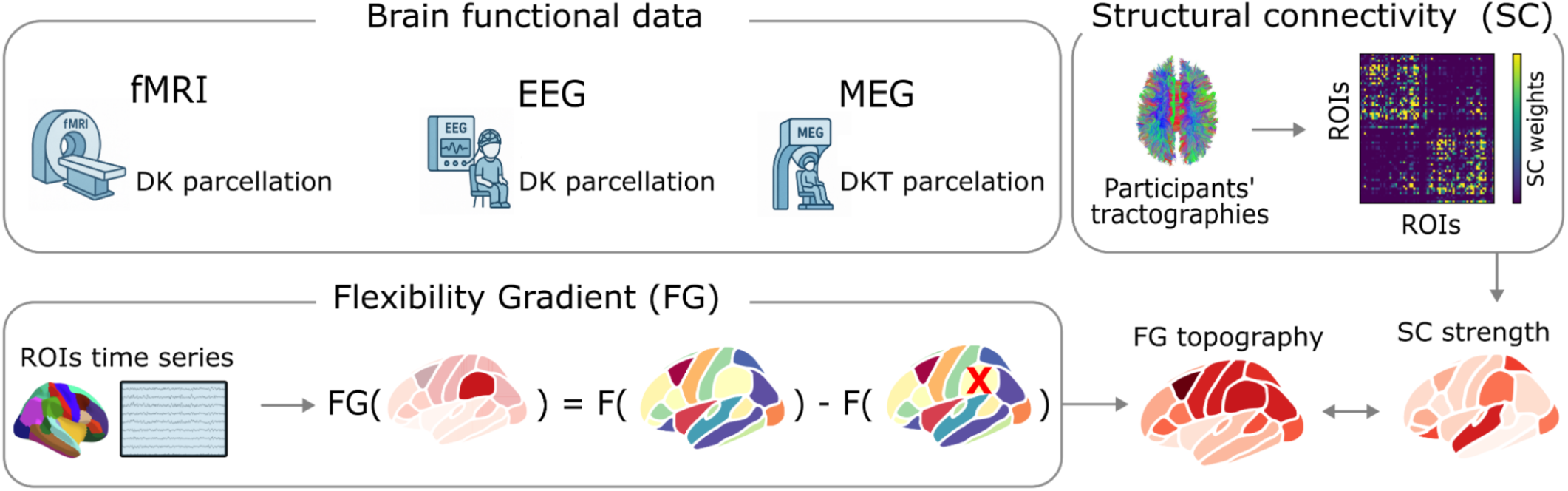
Overview of datasets and flexibility-gradient computation. The top-left panel depicts a schematic representation of the data we employed in our study. In more detail, we investigated fast dynamics using EEG and MEG data and slow dynamics using fMRI data. From the structural connectivity, we quantify centrality by summing the weights of the structural connections within each region, as shown in the top-right panel. Finally, the bottom-left panel presents the definition of the Flexibility Gradient (FG) for region i as the difference between the size of the full functional repertoire and that of the functional repertoire after region i has been removed. The letter F denotes the computation of flexibility.

## RESULTS

To estimate the flexibility of fast brain dynamics, we followed previous work^1–3^. The initial step consists of z-scoring and binning the signal amplitudes of each MEG- and EEG-source-reconstructed activity and the fMRI BOLD time series. At any given time point, a z-score exceeding in absolute value a specified threshold was set to 1 (active), while all other time points were set to 0 (inactive). A burst of activation was defined as a collective phenomenon (i.e., defined at the level of the whole brain), starting when any brain region became active and ending when no region remained active. While bursts of activities are extracted with the same procedure typically used to extract neuronal avalanches, often used to investigate whether brain is operating at criticality, we intentionally avoid this terminology, since our primary aim is to characterize the role of individual brain regions’ flexibility rather than to directly investigate the underlying dynamical regime. For each burst of activity, we extracted the activation patterns as the set of regions active during the burst. *Flexibility* was then quantified as the number of unique activation patterns observed during the recording period, as done in previous literature^2, 9^.

Then, to determine the contribution of each brain region to the overall flexibility, we computed the *flexibility gradient (FG)* for each region R_i_, defined as: FG_i_ = F(R^n^) − F(R^n^_-i_), where F(R^n^) represents the flexibility of all the recorded regions and F(R^n^_-i_) represents the flexibility when region R_i_ is excluded, Figure 1C. For the analysis of fMRI data, we employed the same methodology, with two adjustments to account for the different nature of fMRI measurements (see methods).

### FG heterogeneity across brain lobes

Our results reveal a distinct spatial distribution of the flexibility gradient (FG) across brain regions, with the frontal, parietal, and occipital lobes consistently showing the higher FG values relative to temporal and limbic areas (Fig. 2). Notably, this pattern was present across modalities and temporal scales, in EEG, MEG, and fMRI data, highlighting consistent differences in the way brain regions contribute to the flexibility of brain dynamics. To quantify these differences, we compared FG values across lobes using ANOVA. We accounted for subject and region effects to test for differences across lobes (see methods).

**Figure 2.**
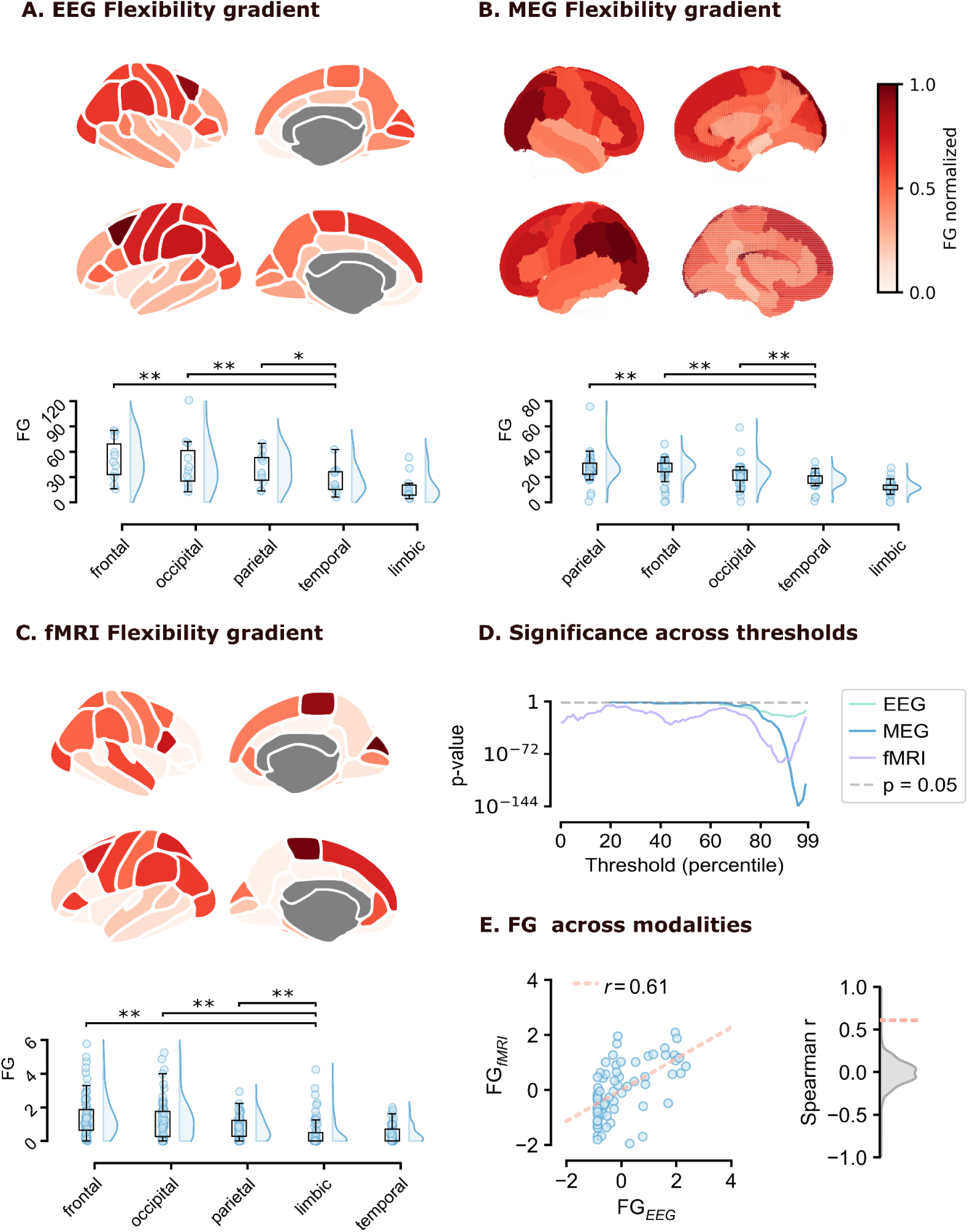
FG topography and invariance over temporal scales. Panels **(A)** and **(B)** show the topography of the FG computed from fast brain dynamics using EEG and MEG data, respectively. Panel **(C)** displays the FG in slow brain activity, computed using fMRI data. EEG and fMRI data were source-reconstructed using the Desikan-Killiany atlas, which segments the cortex into 68 regions. These were grouped into five macro-areas: frontal, parietal, occipital, temporal, and limbic. The distribution of FG across participants in the study is depicted in the lower panels **(A)** and **(C)** for EEG and fMRI data, respectively. For MEG data, panel **(B)**, we used the Desikan-Killiany-Tourville (DKT) atlas, grouping its 84 regions into seven broader areas: frontal, parietal, occipital, temporal, cerebellum, limbic, and subcortical. The distribution of FG across 47 participants is shown for each area in the lower part of panel **(B)**. In panel **(D)**, the p-values from the ANOVA test are largely significant for the thresholds considered, accounting for the mixed effects of brain regions and participants. Panel **(E)** presents a scatter plot comparing the z-score of regional FG values between fMRI (x-axis) and EEG (y-axis), with each dot representing a brain region.

We found significant differences in all datasets (Fig. 2), supporting the existence of consistent heterogeneity in local contributions to global brain flexibility, regardless of the spatial and temporal resolutions of the techniques. Across a broad range of thresholds, the ANOVA yielded significant p-values, around the 50th percentile for fMRI, and above the 75th percentile for MEG and EEG, highlighting the role of high-amplitude bursts of activity in MEG and EEG (Fig. 2D). After identifying significant differences across lobes, we performed post-hoc analysis to evaluate pairwise differences, applying the Holm–Bonferroni correction for multiple comparisons. While the frontal, parietal, and occipital lobes consistently showed significantly higher FG values relative to the limbic and temporal lobes (p≤0.001 for all comparisons between frontal, parietal, or occipital and limbic or temporal), the relationships within these two groups differed across datasets, particularly between fMRI and MEG/EEG. These findings remain robust across threshold choices, in particular for thresholds around the 50th percentile for fMRI and above the 75th percentile for EEG and MEG (Fig. S1A).

Finally, to test the similarity across techniques with different temporal resolutions, we computed the correlation between the FG_fMRI_ and the FG_EEG_, finding a significant correlation between the fMRI and EEG topographies (r(FG_fMRI_, FG_EEG_) = 0.61, p-value≤0.001). This result suggests that the observed differences are robust across temporal scales, spatial resolutions, and recording techniques. This result remains robust across different threshold choices; specifically, EEG thresholds above the 75th percentile and fMRI thresholds around the 50th percentile (Fig. S1B).

### Hemispheric symmetry and correlation with structural connectivity

To better understand the heterogeneity ⥶of local contributions to flexibility, we quantified the hemispheric symmetry of FG topography and its relationship with structural connectivity. The analysis in this section rely on Spearman’s correlations computed (i) between the two hemispheres, to assess hemispheric symmetry, and (ii) between FG and the structural centrality, computed for each brain region as the sum of its structural connections, to examine the contribution of structural connectivity to shaping FG. The significance of these correlations is evaluated using three complementary approaches. First, we compare the observed correlations against a *cyclic-permutation null model*, thereby disrupting spatial organisation while preserving the distribution of values (see Methods). This test evaluates whether empirical correlations exceed those expected under a spatially random configuration. Second, we compute correlations at the individual-subject level and assess whether their population mean is significantly greater than zero using a one-sample *t*-test, thereby quantifying inter-subject consistency. Third, we use a bootstrap procedure to evaluate the stability of the correlations under resampling. To systematically evaluate (i) the robustness of our findings and (ii) by which kind of neural events these results are driven, we systematically tested 99 different threshold values, spanning from 1st percentile of the data distribution to the 99th percentile. Additional details and results are provided in the Supplementary Information. In the main text, we report the threshold ranges for which all three statistical procedures yield significant p-values (see Methods).

Firstly, we tested for hemispheric symmetry. To this end, we computed the correlation between the FG of the left and right hemispheres, finding significant correlation for all three different modalities (MEG: corr(FG_left_, FG_right_) = 0.91, p-values≤0.001 for all the different statistical tests; EEG: corr(FG_left_, FG_right_) = 0.96, p-values≤0.001 for all the different statistical tests; fMRI: corr(FG_left_, FG_right_) = 0.68, p-values≤0.001 for all the different statistical tests). While we reported only the results corresponding to the thresholds maximising the correlation value (MEG threshold = 94th percentile; EEG threshold: 97th; fMRI threshold: 51st), our findings are stable across a broad range of thresholds (all thresholds>25^th^ percentile for fMRI, thresholds>75^th^ percentile for EEG and MEG; Fig. S2 - S3 - S4 - S5 - S6 show in more detail how our results change as a function of the chosen threshold).

Then, we investigated the relationship between the structural connectedness of brain regions and their FG values, following the hypothesis that the number and strength of connections within a brain region contribute to brain flexibility. To test this, we computed the correlation between the FG topography and the strength centrality of brain regions, defined as the sum of all connection weights for a given brain region in the structural connectome. In the structural connectome, the weight of a link corresponds to the number of streamlines connecting the two regions. We found significant correlation for all three different modalities (MEG: corr(SC, FG) = 0.53, p-values≤0.001 for all the different statistical tests; EEG: corr(SC, FG) = 0.56, p-values≤0.001 for all the different statistical tests; fMRI: corr(SC, FG) = 0.45, p-values≤0.001 for all the different statistical tests). While, we reported only the results corresponding to the thresholds maximising the correlation value (MEG threshold = 98th percentile; EEG threshold: 98th percentile; fMRI threshold: 40th percentile), our findings are stable across a broad range of thresholds (20^th^ ≤ threshold≤55^th^ percentile, for fMRI, threshold>80^th^ percentile for EEG and MEG; Fig. S2 - S3 - S4 - S5 - S6 show in more detail the dependence on the threshold of our results).

Importantly, we controlled for the possibility that the FG of brain regions might be trivially driven by the number of above-threshold activations. One could argue that regions that exhibit more activations (i.e., more ones in the binarized time series) are more likely to contribute to unique patterns. As such, the flexibility gradient would not constitute a genuine whole-brain account of flexibility, since univariate regional dynamics would drive changes in the FG. To assess this potential confound, we quantified the correlation between the number of activations and the FG across regions, for all thresholds for the three modalities. For all thresholds used in our main analysis, this correlation was not significant in MEG, EEG, or fMRI, indicating that FG is not simply explained by activation counts. For fMRI, however, correlations became significant at thresholds below 40% and above 60%. For this reason, we restricted the fMRI analysis to the 40–60% threshold range, where activation counts do not trivially account for FG. We also note that, at very high thresholds, the fMRI FG–activation correlation becomes negative, an inversion driven by systematic variations in the number of activations across regions (Fig. S12, S15, and S16).

## DISCUSSION

In this study, we developed a novel framework to investigate how individual brain regions contribute to the flexibility of large-scale neural dynamics across different temporal scales. Central to our paper is the introduction of the *Flexibility Gradient* (FG), a new metric that quantifies the reduction in the size of the dynamic repertoire following the removal of specific brain regions. By applying this method to data recorded using EEG and MEG, which are sensitive to fast electrophysiological dynamics, and fMRI, which are sensitive to slower hemodynamic fluctuations, we were able to characterize the FG and its relationship with brain structural connectivity across a broad temporal spectrum.

Our results reveal a consistent flexibility gradient topography, in which parietal, occipital, and frontal regions contributed most to flexibility. This conclusion is supported by extensive literature on hierarchical modularity and connector-hub architecture, which consistently identifies overlapping community structures with association cortical hubs linking sensory and transmodal areas^16–18^. We find the FG to be symmetric across hemispheres, with homotopic regions showing comparable values, a pattern reminiscent of those classically described in interhemispheric homotopic functional connectivity studies^13,15,19–23^. Interestingly, our results elevate the occipital cortex, traditionally viewed as a sensory and unimodal region, to a primary driver of flexibility, reflecting graph-theoretical evidence that occipital modules occupy a high position in brain hierarchy^24,25^. Notably, this finding does not contradict the unimodal positioning of visual cortex along the functional gradient and sensory-association axis^17,26^, but rather reflects its prominent placement along secondary gradients of cortical organization, where occipital regions occupy an extreme position^17^. This dissociation suggests that regional flexibility is not solely determined by the unimodal–transmodal axis captured by the sensory-motor association axis, but instead emerges from a multidimensional hierarchical architecture in which visual areas play a key dynamic role.

**Table 1.**
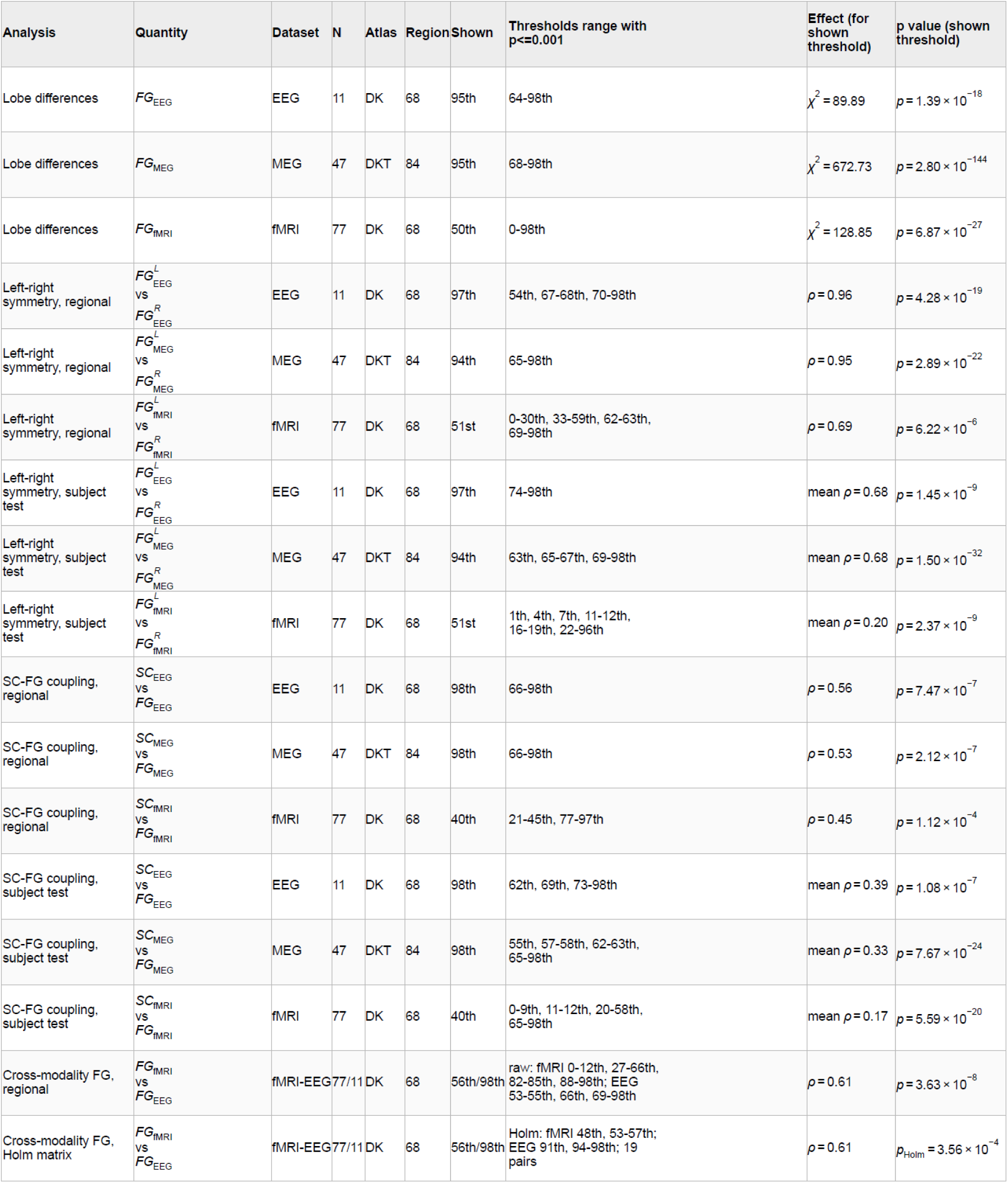
Summary of the main statistical analyses performed across EEG, MEG, and fMRI datasets. For each result, the table reports the modality-specific flexibility gradient measure, sample size, atlas, number of regions, representative threshold shown in the figures, threshold range showing statistical support at p < 0.001, effect size, and statistical significance. FG_EEG_, FG_MEG_, and FG_fMRI_ denote modality-specific flexibility gradients; SC-FG rows quantify structure-function coupling; left-right rows quantify hemispheric symmetry; and cross-modality rows quantify spatial correspondence between FG_fMRI_ and FG_EEG_. For fMRI analyses, threshold-dependent results should be interpreted primarily within the 40th–60th percentile range, where the binarization level was selected as the most physiologically meaningful and comparable regime.

Moreover, the observed FG topography relates to previous work showing that flexibility and temporal variability in brain networks are not uniformly expressed across regions^27–30^. Resting-state studies of dynamic functional connectivity have shown that the connectivity profile of a given region can vary across time windows, and that some brain areas, for instance frontoparietal regions, show stronger temporal variability or more dynamic participation in multiple networks than others^27,28^. In parallel, task-based work on flexible hubs has shown that frontoparietal and cognitive-control regions can rapidly update their whole-brain connectivity patterns across task demands, supporting adaptive control and flexible behaviour^5,31^. Our findings are broadly consistent with this view, as they show that regional contributions to whole-brain flexibility are heterogeneous and particularly strong in frontal and parietal regions. This convergence suggests that the regions contributing most to FG may partially overlap with systems previously implicated in dynamic network reconfiguration and adaptive cognitive control. At the same time, our results also show strong contributions from occipital regions, indicating that FG does not simply reproduce the canonical topography of frontoparietal flexible hubs or resting-state temporal variability.

The FG spatial topography that we found is robust across recording modalities, reinforcing the idea that the ability of brain regions to support flexible dynamics is both anatomically grounded and functionally conserved across different techniques and temporal resolution. In this sense, our work represents an integrative approach in which a research question is investigated using different techniques and systematically across spatial and temporal scales. Following the work of Sejnowsky et al., this is an example of cross-scale and cross-technique integration^32,33^. We tested whether the flexibility gradient (FG) of brain regions was driven by the number of above-threshold activations, under the hypothesis that regions activating more frequently might trivially exhibit higher FG values. This was not the case, except for the fMRI data within a restricted range of thresholds, which we therefore excluded from subsequent analysis.

Across modalities, we established the reproducibility and robustness of our findings through three complementary statistical analyses. First, we performed a permutation-based correlation test that preserves the marginal structure of the data, providing a rigorous benchmark for assessing whether the observed cross-modal correspondence could arise by chance. Second, we applied a bootstrap procedure, demonstrating that the FG topography remains stable across resampled datasets and thus confirming the robustness of our results. Third, we examined reproducibility at the single-subject level, showing that subject-wise FG correlations are consistently and significantly greater than zero, indicating that the observed topographies do not merely reflect group-level aggregation but represent reliable individual patterns. Through these analyses, we assessed the robustness of our results and found that significant effects persisted across a broad range of thresholds. We observed convergent evidence for the stability and reproducibility of the flexibility gradient, reinforcing its validity as a consistent property of large-scale brain dynamics.

Importantly, while our findings are consistent across modalities, there is a clear difference in the binarization threshold between techniques with high temporal resolution (MEG and EEG) and fMRI. In greater detail, our findings regarding MEG and EEG highlight the role of rare, high-amplitude bursts in shaping both the cortical heterogeneity of contribution to brain flexibility and the structure-function coupling. Building on recent evidence suggesting that transient, aperiodic bursts hold functional and clinical significance, we varied the threshold used to define these events^1,6,9,10,34,35^. We found that the relationship between structural connectivity and FG emerged at higher thresholds, where only the most prominent bursts were retained. Regions with higher structural centrality exert greater functional influence across the network during these high-impact moments. This pattern was consistent across EEG and MEG, suggesting that flexible inter-regional communication at fast temporal scales is primarily driven by nonlinear, intermittent peaks rather than continuous oscillatory activities.

In the fMRI data, we observed that at higher thresholds, the flexibility gradient (FG) values were trivially driven by the number of above-threshold events. This effect is likely related to the limited number of recording points and the statistical properties of the fMRI signal. First, fMRI BOLD time series are often modeled as approximately Gaussian in many analytic frameworks (e.g., in general linear modeling and related statistical tests). In this context, fluctuations around the mean might be more informative than rare peaks of activity, which can amplify trivial effects. Second, the relatively small number of time samples in typical fMRI acquisitions can make the number of activation points a dominant factor in FG measures. Finally, the methodological differences in activation event definition between modalities may contribute: in MEG and EEG, single above-threshold events were grouped into collective bursts of activity for the computation of flexibility (see Methods), whereas this grouping was not feasible for the fMRI data due to the smaller number of time points and temporal resolution.

Although our study integrates three complementary neuroimaging modalities and demonstrates reproducibility across spatial, temporal, and methodological dimensions, we acknowledge limitations. In particular, the choice of atlas and preprocessing pipeline may influence regional estimates, and future work should systematically evaluate the impact of parcellation schemes, spatial projections, and signal extraction procedures. Additional validation across distinct spatial and temporal scales, for instance, using neuronal cultures, microcircuit recordings, or in silico experiments, would further strengthen the generality of our findings. Future work, in order to build a general understanding of flexibility across different views, should consider relating our approach grounded in a focus on burst of activity, to other conceptualizations as the ones mentioned above investigating network flexibility at rest or during cognitive tasks^5,27–31^. Recent literature highlighted the role of higher-order interactions in shaping brain dynamics^36–41^; a key direction for future research will be to determine how groups of regions jointly contribute to brain flexibility and whether specific subnetworks exhibit synergistic or cooperative patterns. Finally, developing mechanistic models to account for the emergence of the Flexibility Gradient could offer a simple yet insightful approach to describe whole-brain dynamics in terms of overall flexibility.

Altogether, our findings put forward a new spatiotemporal map of regional influence on whole-brain dynamics. The Flexibility Gradient integrates fast and slow modalities to reveal how specific brain regions, especially those in the parietal and frontal cortices, serve as flexible hubs that orchestrate complex dynamics over time and space. This multimodal approach opens new avenues for studying brain function in health and disease, particularly in conditions where flexibility and reconfiguration are impaired.

## METHODS

### Estimation of flexibility

Throughout our analysis, a central role is played by the computations of flexibility, which we perform as previously done in the literature. To this end, we first binarized the absolute value of brain data using a threshold computed as the n^th^ percentile, as in previous work. Then, we identify a burst of activity as a set of continuous above-threshold activations: a burst starts when at least one region is above threshold and ends when no regions are above threshold. For each burst of activity, we identify an activation pattern: the set of regions that presented at least an event above threshold during the burst. The flexibility is defined as the number of unique patterns (defined across all the brain regions) observed during the recording.

For the analysis of fMRI data, we employed the same methodology, with two adjustments to account for the different nature of fMRI measurements. First, binning was not performed based on absolute signal values, because low BOLD amplitudes represent periods of low activity, whereas in EEG and MEG, both positive and negative peaks may correspond to abrupt electrophysiological events. Second, given the low temporal resolution of fMRI, each time point was treated as an activation pattern, and flexibility was computed as the number of unique activation patterns recorded. This choice was motivated by the relatively short duration of the fMRI recordings (242 time points), which does not permit the reliable grouping of consecutive activations into bursts. All other steps for computing the FG were kept identical across modalities.

### Estimation of FG

For each brain region r_i_, its flexibility gradient, FG(r_i_), is defined as the difference between the flexibility of the whole brain and when excluding the region r_i_. To assess the influence of region ri on the number of patterns, we stored the unique patterns and computed the reduction in the number of unique patterns upon subtracting region r_i_. This way, we observed directly the influence of brain regions on the number of unique patterns explored.

### fMRI data set

The fMRI data were taken from the eBRAINS public dataset “Structural and functional connectivity data of the ARCHI database in the Desikan atlas (v1)” (DOI:10.25493/91BN-SZ9), derived from the CONNECT/ARCHI project^42,43^. The original cohort consists of healthy young adults scanned at NeuroSpin (CEA, France) on a 3T Siemens Tim Trio with a 12-channel head coil. The resting-state fMRI acquisition comprised a 10-min gradient-echo EPI sequence (40 axial slices, 3 mm isotropic voxels, TR = 2.4 s, TE = 30 ms, flip angle = 90°, ascending interleaved slice order). From this dataset on eBRAINS, 77 subjects were publicly available, for whom both functional and diffusion MRI had already been preprocessed.

Structural connectivity data for the same individuals were obtained from the same eBRAINS resource Structural and functional connectivity data of the ARCHI database in the Desikan atlas, which provides diffusion MRI–based connectivity matrices derived from the ARCHI cohort and mapped onto the Desikan–Killiany parcellation. As described by the previous publication of this dataset^43^, diffusion data were processed using the BrainVISA/Connectomist toolbox, and whole-brain deterministic streamline tractography was performed for each subject. The resulting tractography data were used to derive cortical parcels via connectivity-based clustering, and structural connectivity matrices were obtained for each subject and mapped onto the Desikan-Killiany atlas^44^.

### EEG data set

Details regarding EEG preprocessing can be found in the supplementary materials of Schirner et al^45^. Initially, to mitigate slow drifts in EEG signals and enhance the construction of artifact templates during MR imaging artifact (IAA) correction, all EEG channels were high-pass filtered at 1.0 Hz using a standard FIR filter. IAA correction was conducted with Analyser 2.0 software (version 2.0.2.5859, Brain Products, Gilching, Germany). The onset of each MRI volume was identified using a gradient trigger threshold of 300 µV/ms. Markers associated with shimming artifacts or large movements were manually excluded. To ensure accuracy, the inter-scan intervals were checked for consistency with the expected repetition time (TR) of 1940 ms.

For each EEG channel, IAA templates were generated using a sliding average approach with a window of 11 intervals and subsequently subtracted from the corresponding scan intervals. The EEG data were then downsampled to 200 Hz, imported into EEGLAB, and low-pass filtered at 60 Hz. ECG traces were employed to detect QRS complexes and mark the occurrence of ballistocardiogram (BCG) artifacts. The timing and placement of ECG markers were visually inspected and corrected when necessary.

To remove BCG and movement-related artifacts (particularly eye movement), temporal independent component analysis (ICA) was performed using the extended Infomax algorithm in EEGLAB. Identification of independent components (ICs) containing BCG artifacts was based on their topographies, activation time courses, power spectra, and heartbeat-locked averaged potentials. Components identified as BCG-related were excluded from back-projection. Additional artifact-related ICs, mainly associated with movement and eye activity, were recognized by their spatial distribution, temporal characteristics, spectral profiles, and event-related potentials.

Structural connectivity was derived from diffusion-weighted MRI data using a standardized processing pipeline. Following correction for head motion and eddy-current distortions, whole-brain fiber tracking was performed in native diffusion space. Cortical and subcortical regions were defined according to an anatomical parcellation obtained from individual T1-weighted images. Streamlines were assigned to region pairs to construct subject-specific structural connectivity matrices, where connection weights reflected the number (or density) of reconstructed fibers between regions.

The anatomical parcellation was based on the Desikan–Killiany cortical atlas^44^. Comprehensive descriptions of EEG time series and tractography preprocessing are available in prior publications (Becker et al., 2011; Freyer et al., 2009a; Ritter et al., 2007; Ritter et al., 2010)^45–49^.

### MEG data set

The MEG dataset was previously published in Sorrentino et al, 2021^10^. In a nutshell, the data set contains recording of resting state MEG activity of 47 young adults from the general population. MEG recordings were ∼7 minutes long. MEG data preprocessing and source reconstruction were conducted according to the procedures described in Sorrentino et al. (2021)^10^. In short, environmental noise was removed via principal component analysis based on reference sensors. Supervised ICA was used to eliminate physiological artifacts such as eye blinks and cardiac activity. Noisy channels were visually identified and excluded by an expert rater. Neural time series were reconstructed for 84 DKT atlas regions. Source modeling employed Nolte’s (2003)^50^ volume-conduction model and the linearly constrained minimum variance (LCMV) beamformer algorithm^51^, using individual structural MRIs. Source activity was estimated at the centroid of each ROI. All preprocessing and source reconstruction steps were implemented in the Fieldtrip toolbox (Oostenveld et al., 2011). Structural brain images were acquired using a 1.5 Tesla scanner (Signa, GE Healthcare). MRI acquisition followed the MEG recordings.

For tractography analysis, we used ROIs from the MNI-based DKT atlases, masked with the SPM gray matter probability map (threshold = 0.2). FA maps were normalized to MNI space using FSL templates, and spatial normalisation was performed using SPM12. Normalization accuracy was confirmed visually. From each subject’s whole-brain tractography and corresponding GM ROIs, the number of streamlines connecting each ROI pair was computed using in-house software.

### Lobe-level differences

To test whether the mean value of our metric differed across cerebral lobes, we fitted a linear mixed-effects model with lobe as a fixed effect, random intercepts for subjects to account for repeated measures, and a variance component for brain regions nested within lobes to capture region-level variability. Each observation represents the value for subject i, lobe j, and region k. The omnibus null hypothesis was that all lobe means were equal. We compared the full model (including lobe) with a reduced model (excluding lobe) using a likelihood-ratio test under maximum likelihood estimation. All tests were two-sided with alpha = 0.05, and analyses were performed in Python using the statsmodels MixedLM implementation. Missing or invalid values were removed listwise prior to fitting.

When the omnibus test was significant, we conducted pairwise comparisons between lobes using Wald tests on model-based contrasts of the fixed effects. For each pair of lobes A and B, we tested whether the difference in their fixed-effect estimates was zero. Resulting p-values were adjusted across all pairwise tests using the Bonferroni-Holm method to control for multiple comparisons.

### Statistical tests for Spearman correlations

To evaluate the significance and robustness of the Spearman correlations reported in this study, we used three complementary statistical approaches, following standard nonparametric inference procedures commonly used in neuroimaging^52^.

First, we constructed a nonparametric *cyclic-permutation null model*. For each threshold, we generated 1000 permutations and recomputed the Spearman correlation for each permuted dataset. This procedure breaks any spatial or anatomical correspondence between the two variables while preserving their marginal distributions, thereby providing an estimate of the correlation expected under random regional assignment. Permutation p-values were computed as the proportion of null correlations whose absolute value was greater than or equal to the absolute empirical correlation. Because permutations were performed independently at each threshold, no correction for multiple comparisons was applied to these p-values. The minimum attainable p-value with 1000 permutations was therefore 0.001. Results from this null model were consistent with those obtained using the other statistical tests and are shown across different thresholds in Figures S4, S5, S9, and S10.

For each participant, we computed the Spearman correlation between FG and structural connectivity (SC) strength, as well as between FG values in the left and right hemispheres. Note that in all our analysis SC strength centrality is computed from the structural connectivity matrix, for each brain region, as the sum of all the weights of its structural connections. This resulted in one correlation coefficient per subject for each analysis. We then assessed whether these subject-level correlations were consistently greater than zero at the population level. For each threshold used in the binarisation procedure, we performed a one-sample t-test against zero. Because we tested 99 thresholds, p-values were Bonferroni corrected using a factor of 99. In Figure 3, statistical significance is reported using the corrected threshold p = 0.05/99. Results from this test are shown across different thresholds in Figures S2, S3, S7, and S8.

**Figure 3.**
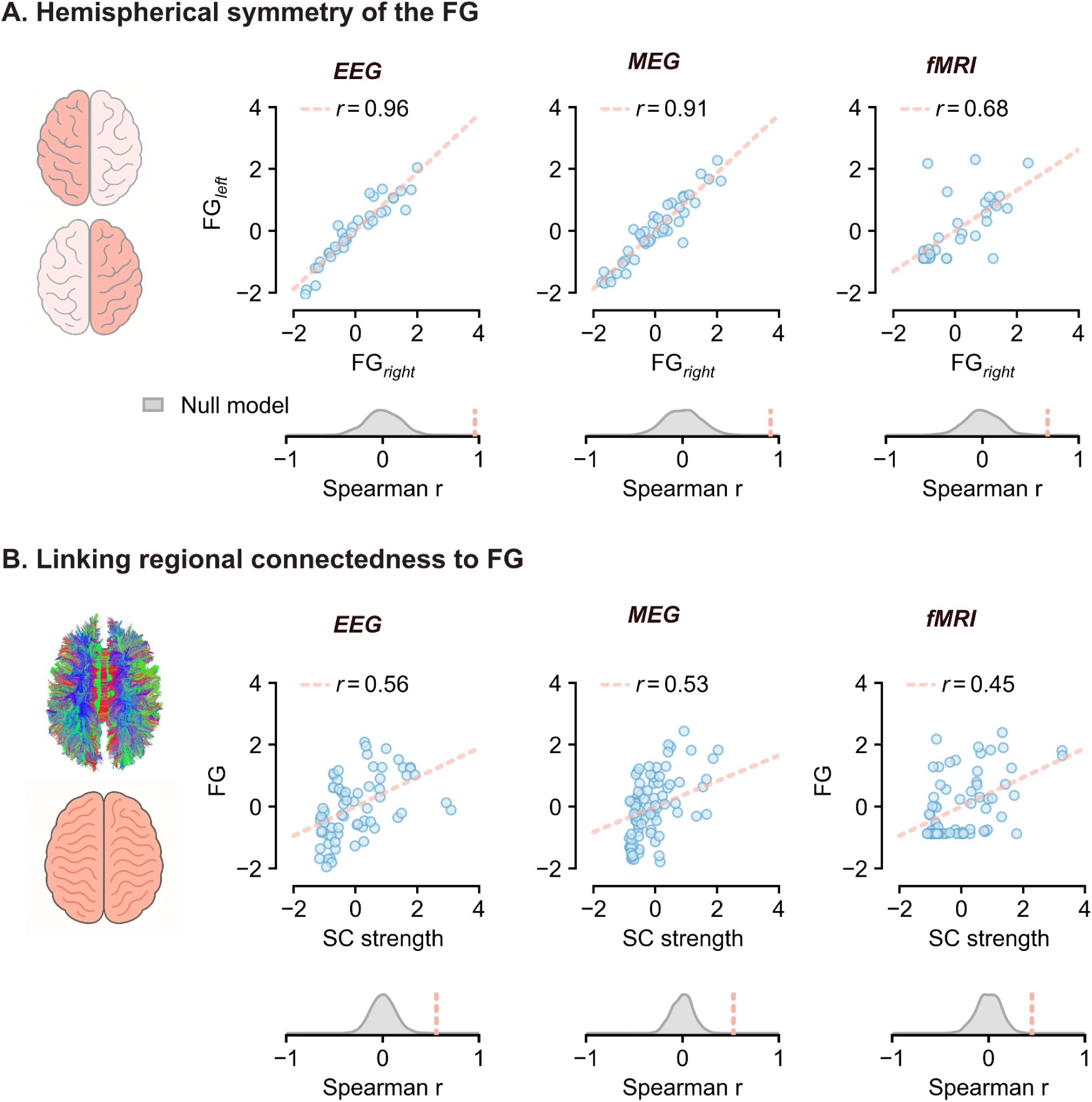
Hemispheric symmetry and structure–function coupling across different temporal scales and data types. This figure illustrates the correlation between structural connectivity (SC) strength-centrality and the FG across brain regions as well as the correlation between the FG of one hemisphere and the other one. In panel **(a)**, upper part, each data point represents a region, with the average FG value across subjects on the x-axis for the left hemisphere and on the y-axis for the right hemisphere. For both axes, values are z-scored before plotting, by subtracting the mean across regions and dividing by the standard deviation. Correlations are computed using subject-averaged FG values and Spearman correlation. In panel **(a)**, lower part, the observed correlation (coral line) is compared to a null distribution (grey) generated by a random shuffle of the data, see methods. In panel **(b)**, upper part, each data point represents a region, with the average FG value across subjects on the x-axis and average SC strength on the y-axis. Both average FG and average SC values are z-scored before plotting, by subtracting the mean across regions and dividing by the standard deviation. Correlations are computed using subject-averaged SC and FG values and Spearman correlation. In panel **(b)**, lower part, the observed correlation (coral line) is compared to a null distribution (grey) generated by a random shuffle of the data, see methods.

Finally, we assessed the stability of our results using a nonparametric bootstrap. For each threshold, we repeatedly resampled brain regions with replacement and recomputed the Spearman correlation on each bootstrap sample (1000 iterations). This approach tests how sensitive the correlation is to the contribution of individual regions by mimicking repeated sampling from the underlying distribution. The resulting bootstrap distribution was used to derive empirical confidence intervals and to assess the robustness of the observed correlations. Bootstrap estimates were highly consistent with the permutation and population-level tests, further supporting the reliability of our findings. Bootstrap confidence intervals (CIs) are shown in figures S6 and S11.

## Data, code, and materials availability

The fMRI resting-state time series of the 77 participants employed in this study and their SC are publicly available through eBRAINS under DOI: https://doi.org/10.25493/91BN-SZ9. MEG and EEG time series and the correspondent SC are already investigated by previous work (for the MEG datasets, DOI: https://doi.org/10.7554/eLife.67400 - for the EEG datasets DOI: https://doi.org/10.1016/j.neuroimage.2015.03.055). All code required to reproduce the main analysis and figures is public on github at the following link: https://github.com/Mattehub/FG.git. Any additional information required to reanalyze the data is available from the corresponding author upon reasonable request.

## Competing interest

The authors have no conflicts of interest.

## Authors contributions

Conceptualization: M.N., M.A. and P.S.; Methodology: M.N., P.S., and M.A.; Software: M.N. and E.B.; Formal analysis: M.N., M.A. and E.B.; Investigation: M.N., M.A., E.B., C.M., E.T.L., M.D., M.D.L., E.G., and M.Q.; Resources: A.B.; Data curation: C.M., E.T.L., M.D., M.D.L., E.G., and M.Q.; Writing - original draft: M.N.; Writing - review and editing: all authors; Visualization: M.N.; Supervision: P.S.

## Funding

M.N. acknowledges the funding he received from the French government under the ‘France 2030’ investment plan managed by the French National Research Agency (Agence Nationale de la Recherche; reference: ANR-16-CONV000X / ANR-17-EURE-0029) and from Excellence Initiative of Aix-Marseille University- A*MIDEX (AMX-19-IET-004).

## Supplementary material

## Threshold exploration for ANOVA and the relationship between fMRI and EEG

In the following sections, we provide a detailed description of the systematic analysis performed to assess the influence of threshold selection on the results reported in the main manuscript. As described in the Methods, we evaluated the significance of lobe-level differences using a linear mixed-effects ANOVA. Figure S1A shows the corresponding post-hoc comparisons, demonstrating that the pattern of lobe differences is highly robust across a wide range of thresholds.

We also examined the significance of the Spearman correlation between FG_fMRI_ and FG_EEG_ using the statistical functions implemented in scipy.stats. As shown in Figure S1B, the correlation remained consistently significant for fMRI thresholds around the 50th percentile and for EEG thresholds above the 75th percentile.

**Figure S1.**
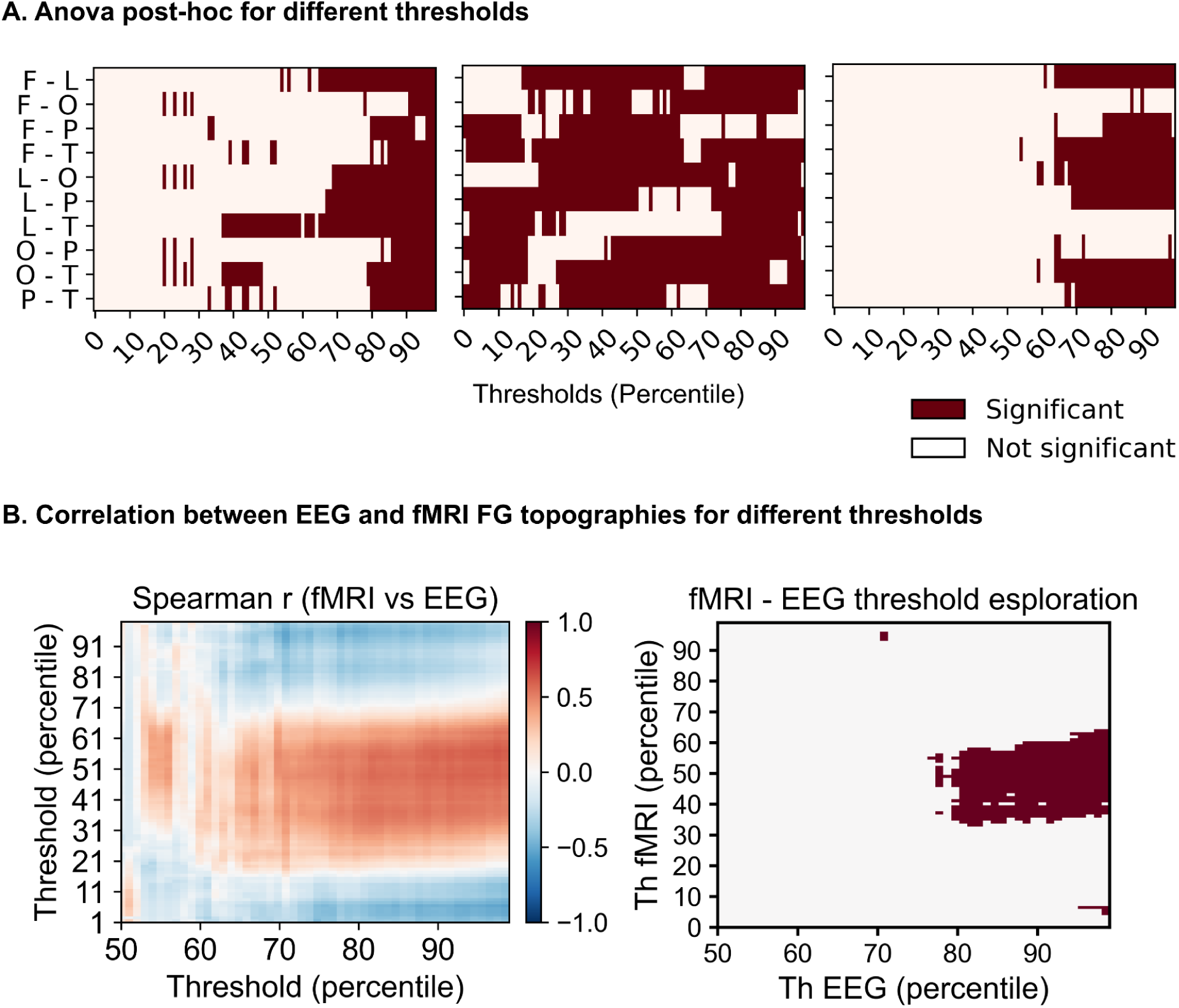
Validity across different thresholds. This figure complements Fig. 2 of the main manuscript by assessing the robustness of the reported results across different threshold values. In panel A, we show the post-hoc ANOVA results for each pair of brain lobes (y-axis) and each threshold (x-axis). Each entry indicates whether the corresponding post-hoc comparison was significant after correction for multiple comparisons using the Holm–Bonferroni method (p < 0.05). Lobe abbreviations are as follows: frontal (F), parietal (P), occipital (O), temporal (T), and limbic (L). For example, the row labelled F–O indicates whether the flexibility-gradient contributions differed significantly between frontal and occipital regions. The mixed-effects ANOVA included brain lobe as a fixed effect, with subject and regions nested within lobes modelled as random effects; further details are provided in the Methods section. Panels B and C show the threshold-dependence of the correlation between EEG-derived and fMRI-derived flexibility-gradient topographies. Because this analysis involves two binarisation thresholds, one for the EEG data and one for the fMRI data, the results are displayed as matrices, with EEG thresholds on the x-axis and fMRI thresholds on the y-axis. In panel B, each matrix entry represents the correlation between the EEG and fMRI flexibility-gradient topographies for a specific pair of EEG and fMRI thresholds. Panel C shows the corresponding statistical significance of these correlations, with significant correlations shown in red and non-significant correlations shown in white.

## Threshold exploration for hemispheric symmetry

In addition, we assessed the robustness of all correlation results using the three statistical pipelines described in the Methods:

1. **Population-level statistics** were used for left-right symmetry analyses, figures S2 and S3, and for structure-function relationships, figures S7 and S8. For each threshold and participant, we computed the correlation between FG_left_ and FG_right_. The distribution of these correlations across participants was then plotted for each threshold. P-values were obtained using a one-sample t-test to assess whether the average correlation was significantly different than zero. The p-values are reported for each threshold in a separate panel, with p = 0.05 highlighted by a red dotted line.
2. **Permutation-based null models** were used for left-right symmetry analyses, figures S4 and S5, and for structure-function relationships, figures S9 and S10. Regional labels were cyclically permuted to estimate the probability of randomly obtaining correlations as large as those observed. For each threshold, the distribution of correlations computed between the averaged FG_left_ and FG_right_ after permutation (n = 1000 iteration) is shown in grey and compared with the observed correlation, indicated by a red star. The corresponding p-values were computed as the proportion of permutations yielding a correlation whose absolute value was greater than or equal to the absolute value of the empirical correlation.
3. **Bootstrap resampling** was used for hemispheric symmetry, figure S6, and structure-function relationships, figure S11. Brain regions were resampled with replacement to quantify the robustness of the correlations across repeated samples. The 95th-percentile confidence intervals are shown as grey shaded areas around the average correlation, plotted in black, across all explored thresholds.

These analyses were performed for all three datasets and yielded consistent results, confirming the robustness of the findings presented in the main text. In the following figures, the correlations are tested across a broad range of thresholds.

**Figure S2.**
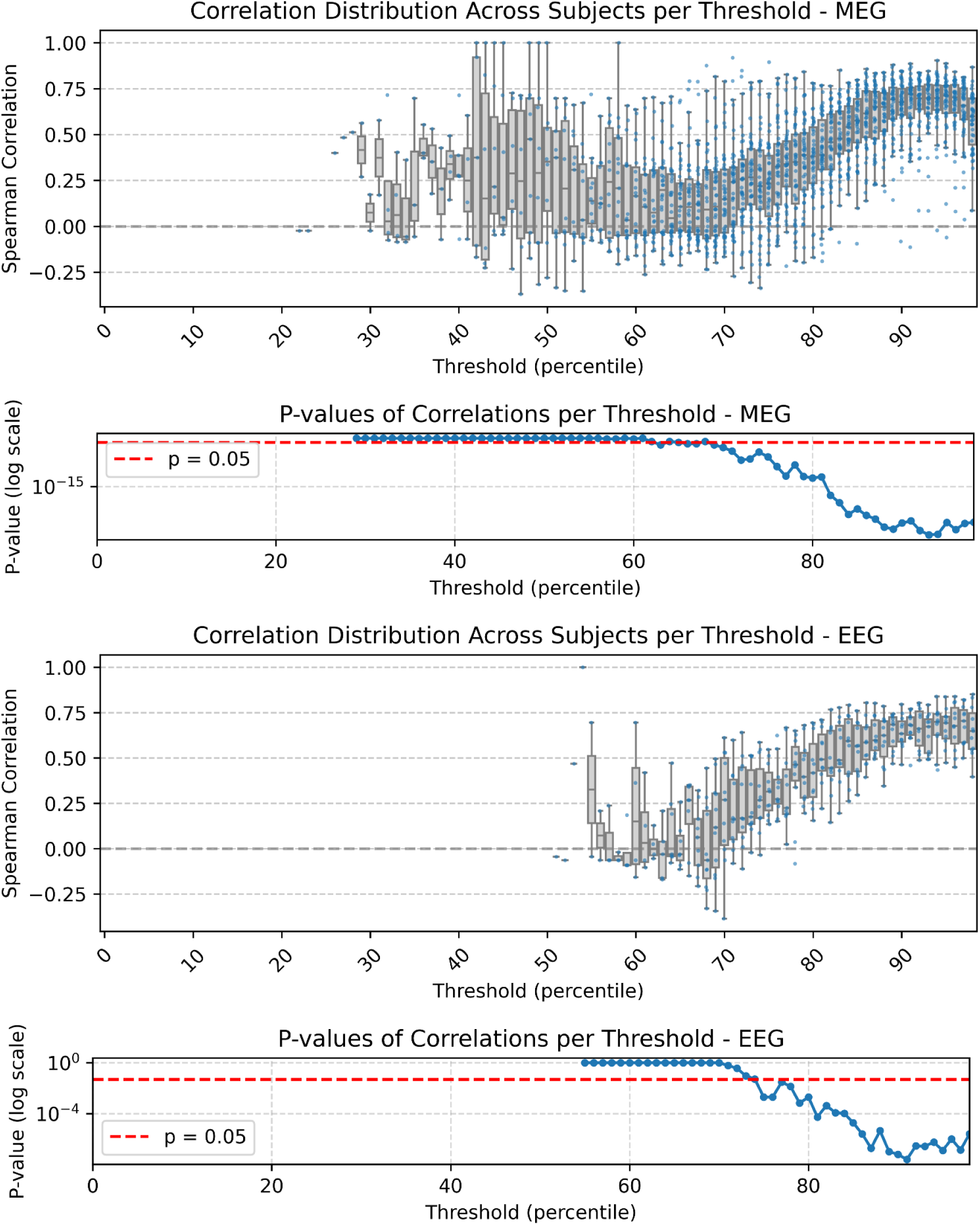
Hemispheric symmetry: validity across different thresholds, one-sample t-test at the population level (EEG - MEG). The figure shows the robustness of hemispheric symmetry in the flexibility gradient (FG) as a function of the binarisation threshold for MEG and EEG data. For each threshold, the Spearman correlation between left- and right-hemisphere FG topographies was computed separately for each participant. The upper panels for each modality show the distribution of subject-level correlations across thresholds, with boxplots summarising the inter-subject variability and individual points indicating participant-specific correlation values. The horizontal dashed grey line marks zero correlation. The lower panels show the corresponding p-values obtained from one-sample tests assessing whether the subject-level Scorrelations were significantly greater than zero at each threshold. P-values are displayed on a logarithmic scale, and the red dashed line indicates the significance threshold of p = 0.05. In both MEG and EEG, hemispheric symmetry becomes stronger and more statistically robust at higher thresholds, particularly above approximately the 75th percentile, indicating that high-amplitude events reveal a consistent bilateral organisation of the flexibility gradient.

**Figure S3.**
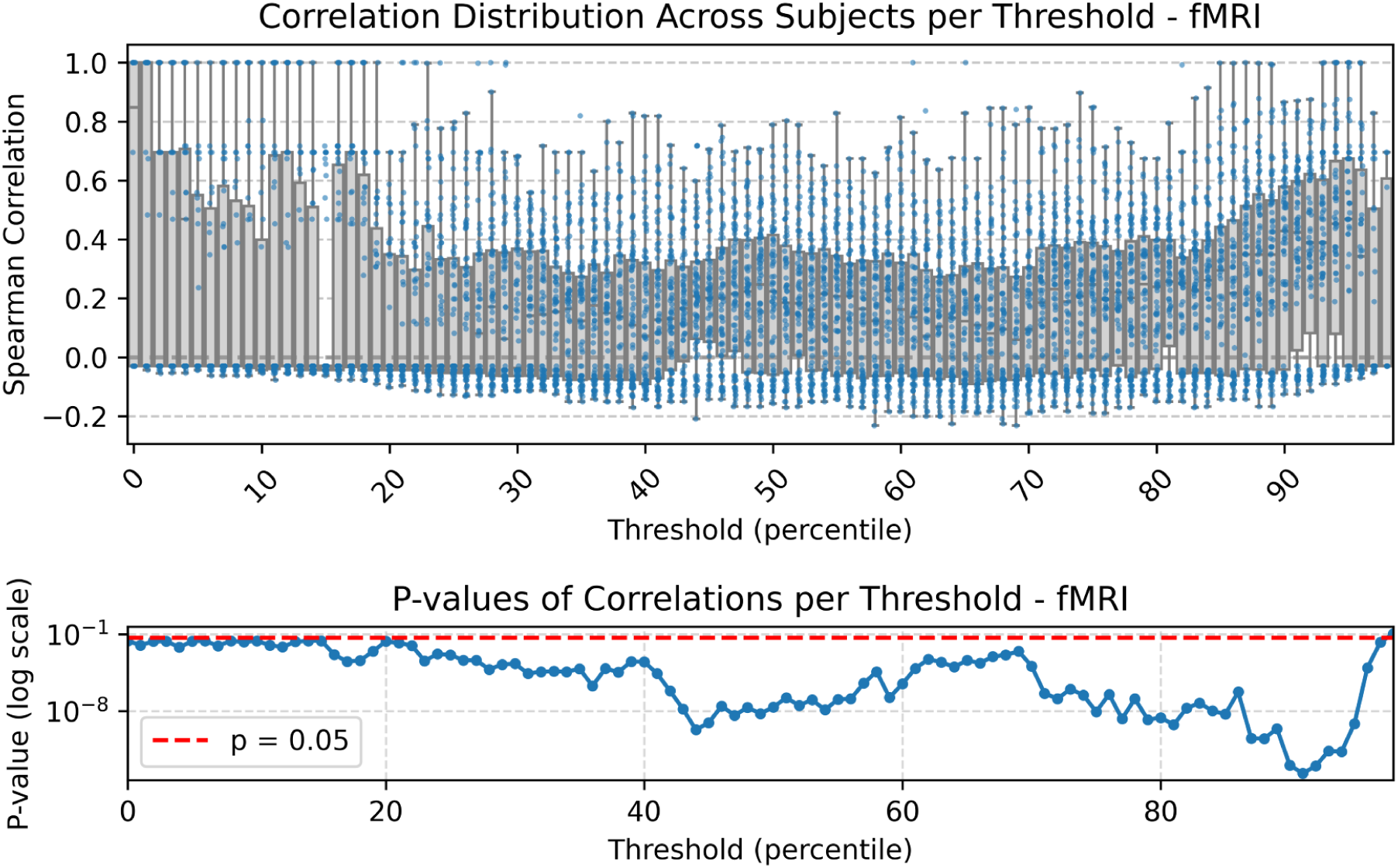
Hemispheric symmetry: validity across different thresholds, one-sample t-test at the population level (fMRI). Same analysis as shown in Figure S2 for MEG and EEG, but applied to the fMRI dataset. For each threshold, the Spearman correlation between the flexibility-gradient topographies of the left and right hemispheres was computed separately for each subject. The upper panel shows the distribution of subject-level correlations across thresholds, while the lower panel reports the corresponding p-values from a one-sample test against zero. P-values are shown on a logarithmic scale, and the red dashed line indicates p = 0.05.

**Figure S4.**
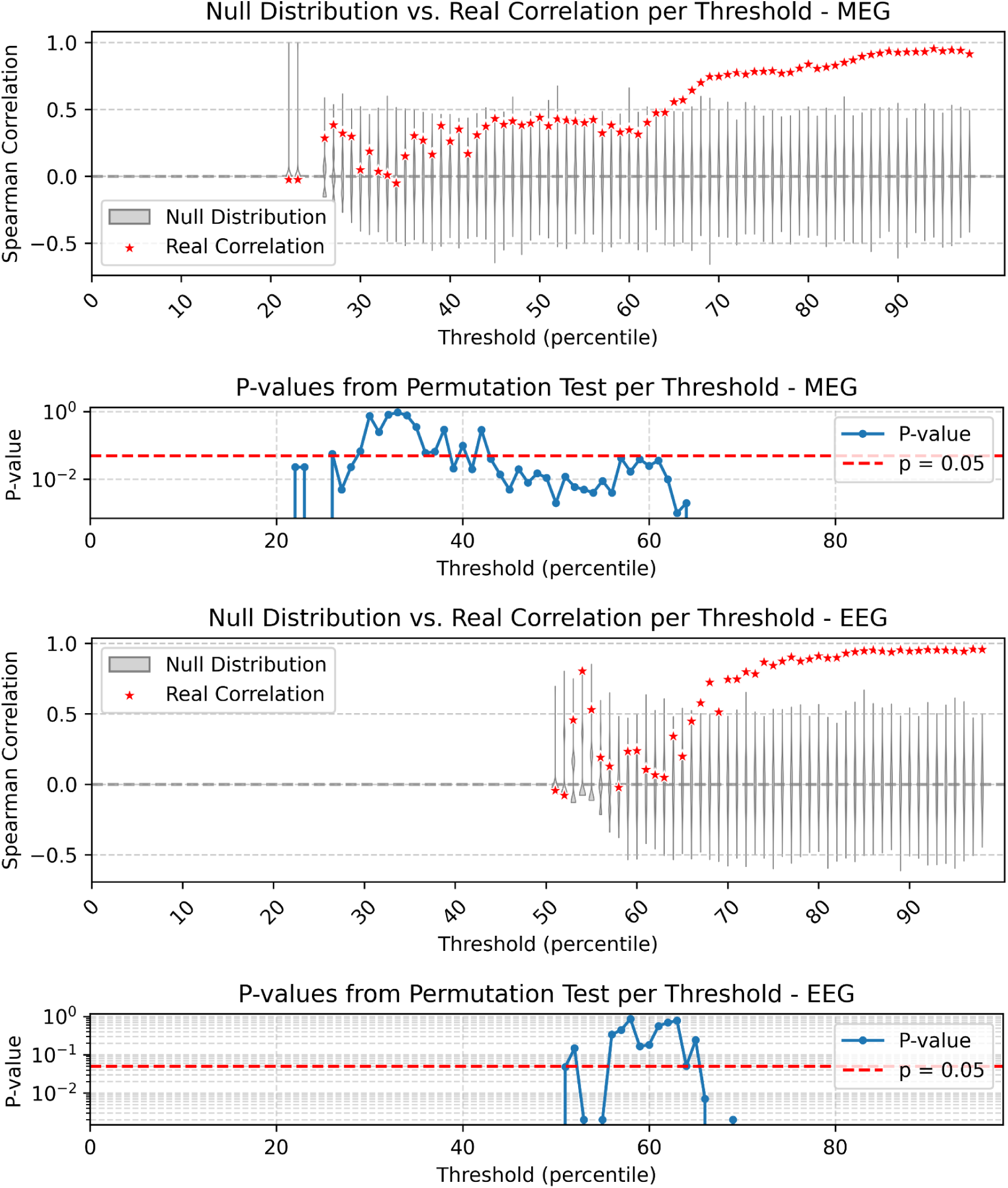
Hemispheric symmetry: validity across different thresholds, null model (EEG - MEG). This figure evaluates the robustness of the hemispheric symmetry of the flexibility gradient (FG) in MEG and EEG data using a cyclic-permutation null model. For each threshold, the empirical Spearman correlation between left-and right-hemisphere FG topographies is shown with red stars. The grey distributions represent the null correlations (n = 1000 permutations) obtained after permuting the regional data before recomputing the correlation, thereby preserving the structure of the data while disrupting the original left–right correspondence (see methods). The lower panels show the corresponding permutation-test p-values across thresholds, with the red dashed line indicating the significance threshold of p = 0.05. In both MEG and EEG, the empirical correlations exceed the null distribution at higher thresholds, supporting a non-random hemispheric symmetry of the FG.

**Figure S5.**
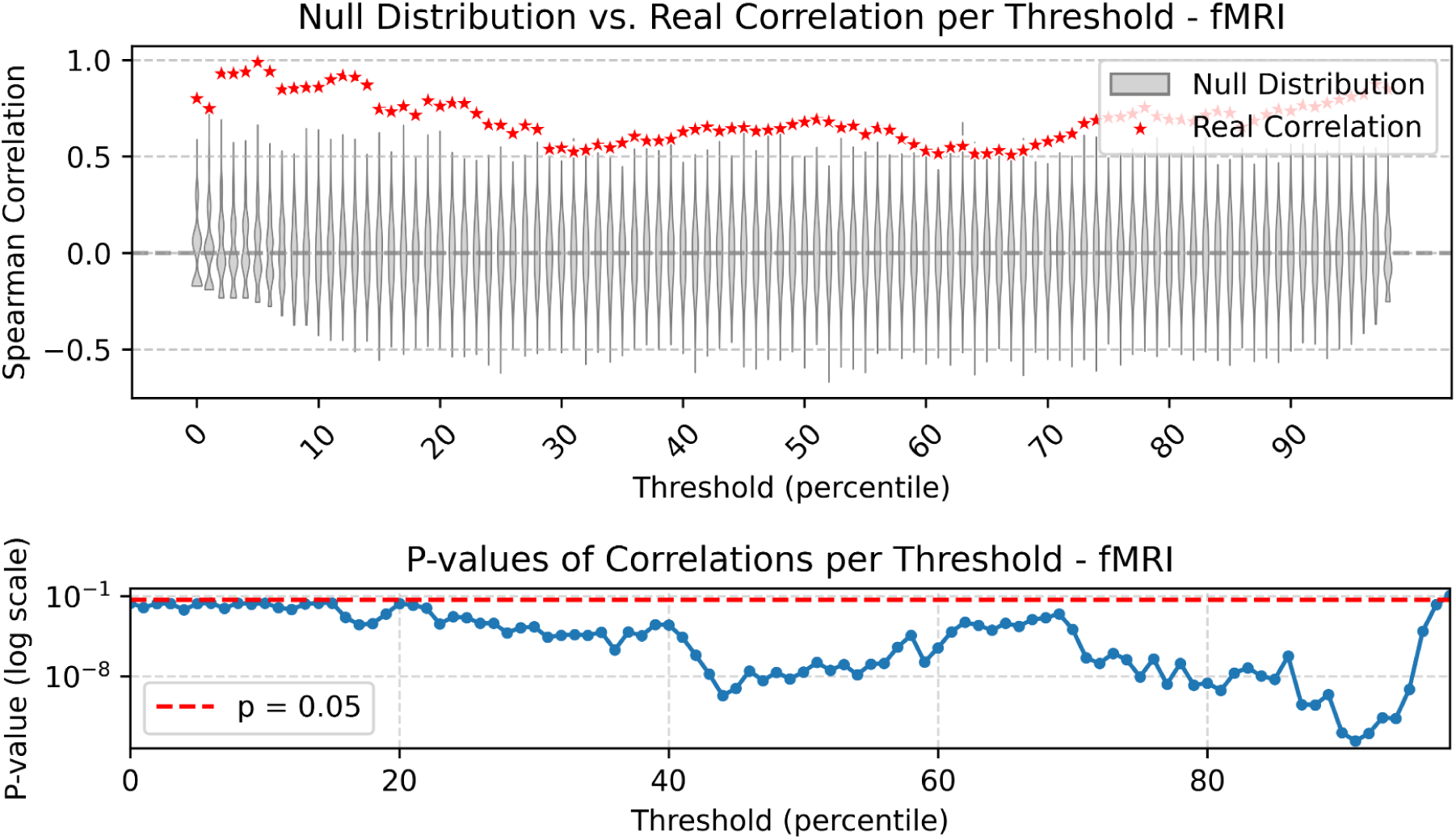
Hemispheric symmetry: validity across different thresholds, null model (fMRI). Same analysis as in Figure S4 for MEG and EEG, but applied to the fMRI dataset. For each threshold, the empirical Spearman correlation between left- and right-hemisphere flexibility-gradient topographies is shown with red stars. The grey distributions represent the null correlations obtained after cyclically permuting the regional data before recomputing the correlation. The lower panel shows the corresponding permutation-test p-values across thresholds, displayed on a logarithmic scale. The red dashed line indicates the significance threshold of p = 0.05.

**Figure S6.**
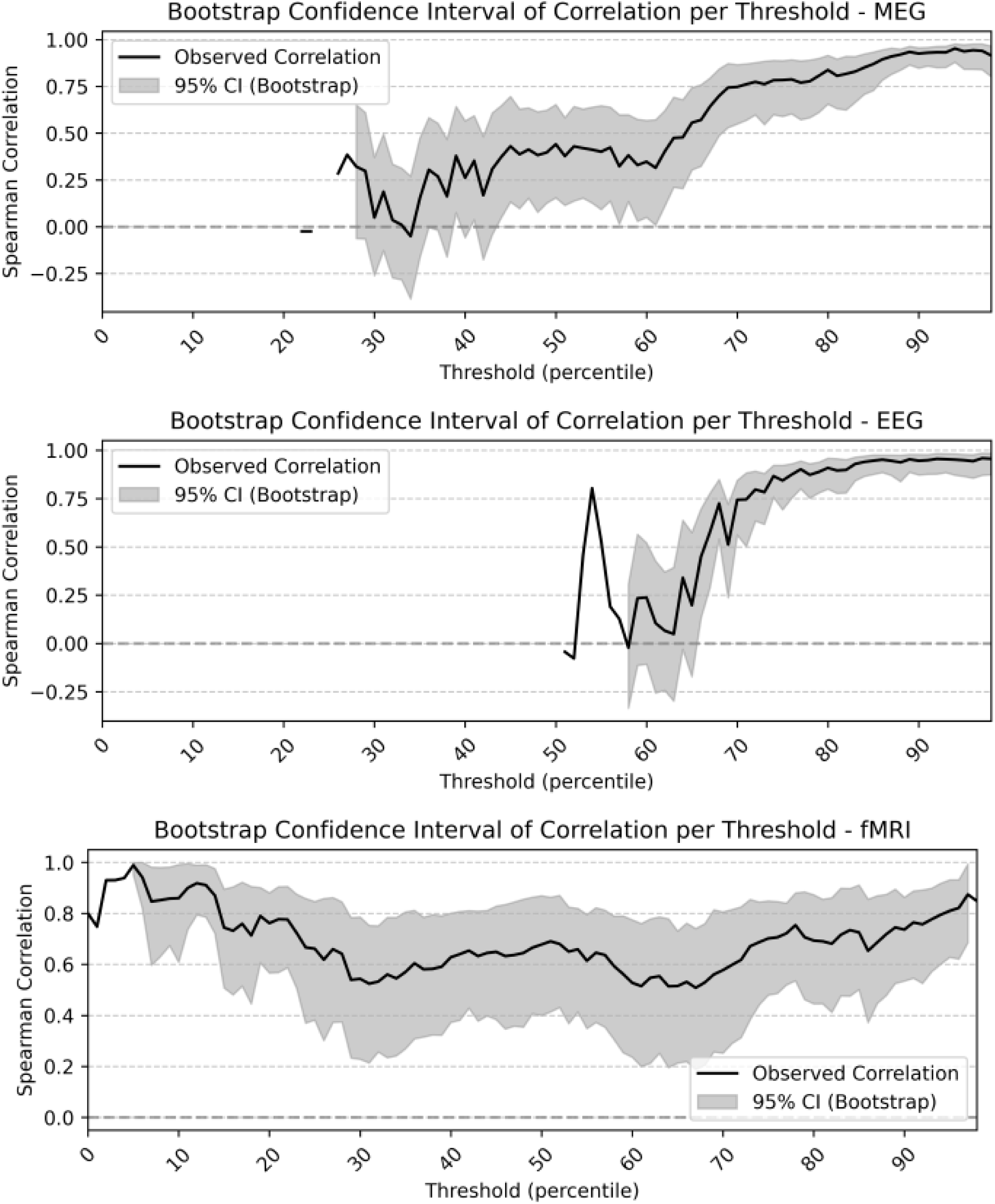
Hemispheric symmetry: validity across different thresholds, bootstrap. This figure shows the stability of the hemispheric symmetry of the flexibility gradient (FG) across different binarisation thresholds for MEG, EEG, and fMRI data. For each threshold, the observed Spearman correlation between left- and right-hemisphere FG topographies is shown as a black line. The grey shaded area represents the 95% confidence interval obtained by bootstrap resampling. Across the three modalities, the correlations remain positive over a broad range of thresholds, with stronger and more stable hemispheric symmetry emerging at higher thresholds for MEG and EEG. For fMRI, the hemispheric correlation remains consistently positive across nearly the entire threshold range.

## Threshold exploration for structure-function correlations

In addition, we assessed the robustness of the correlation between SC strength and FG using the three statistical pipelines described in the Methods for a broad range of different thresholds:

4. **Population-level statistics** (figures S7 and S8). For each threshold and participant, we computed the correlation between FG and SC strength. The distribution of these correlations across participants was then plotted for each threshold. P-values were obtained using a one-sample t-test to assess whether the average correlation was significantly different than zero. The p-values are reported for each threshold in a separate panel, with p = 0.05 highlighted by a red dotted line.
5. **Permutation-based null models** (figures S9 and S10). Regional labels were randomly cyclically permuted to estimate the probability of randomly obtaining correlations as large as those observed. For each threshold, the distribution of correlations computed between the averaged FG and the average SC strength after permutation (n = 1000 iterations) is shown in grey and compared with the observed correlation, indicated by a red star. The corresponding p-values were computed as the proportion of permutations yielding a correlation whose absolute value was greater than or equal to the absolute value of the empirical correlation.
6. **Bootstrap resampling** (figure S11). Brain regions were resampled with replacement to quantify the robustness of the correlations across repeated samples. The 95th-percentile confidence intervals are shown as grey shaded areas around the average correlation, plotted in black, across all explored thresholds.

These analyses were performed for all three datasets and yielded consistent results, confirming the robustness of the findings presented in the main text.

**Figure S7.**
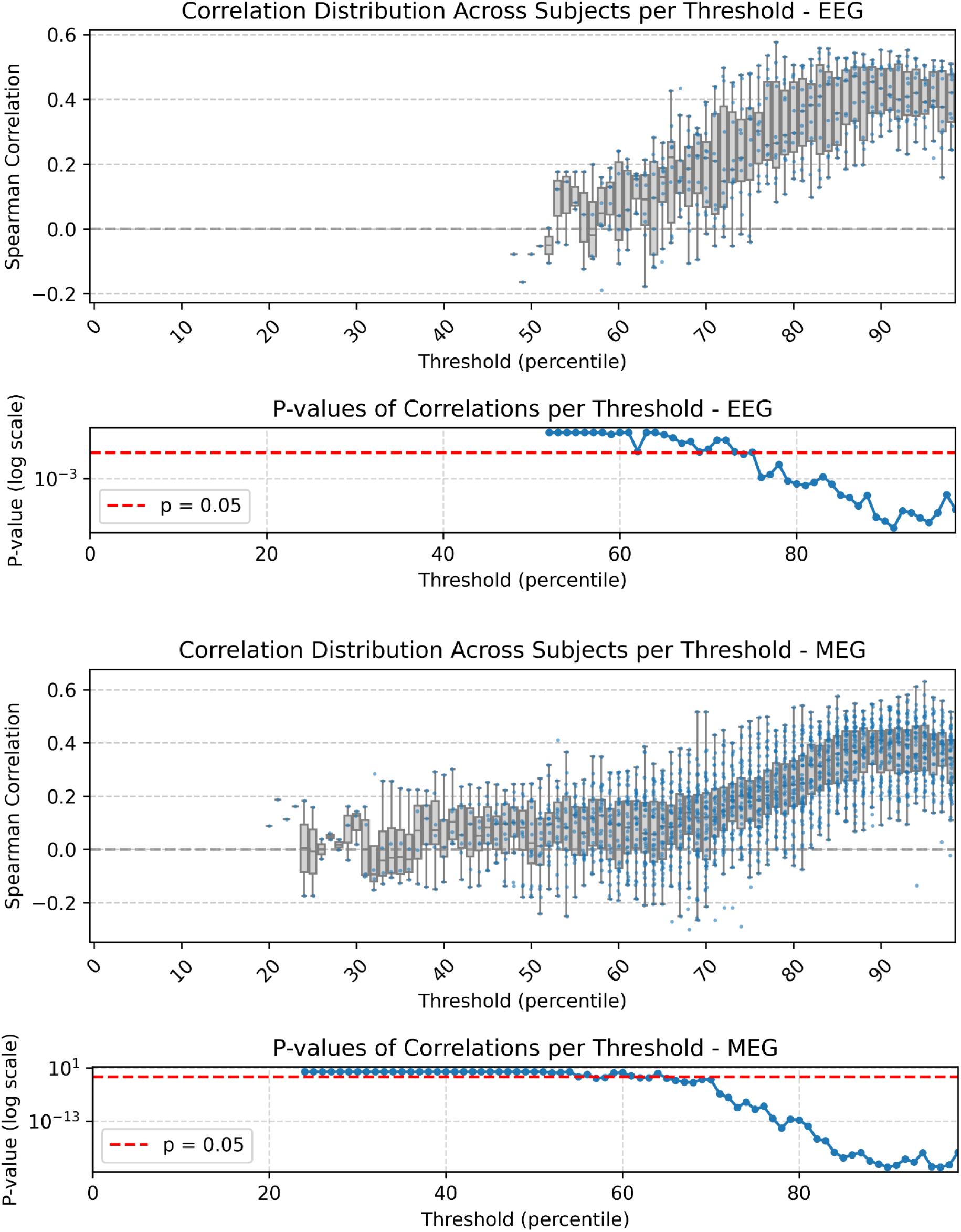
Structure-function correlation: validity across different thresholds, one-sample t-test at the population level (EEG - MEG). This figure shows the robustness of the relationship between structural connectivity (SC) strength and the flexibility gradient (FG) across different binarisation thresholds for MEG and EEG data. This analysis is performed to test the behavior across a broad range of threshold of the correlation reported in Fig.3. For each threshold, the Spearman correlation between regional FG values and regional structural strength was computed separately for each subject. The upper panels for each modality show the distribution of subject-level correlations across thresholds, with boxplots summarising inter-subject variability and individual points indicating participant-specific correlation values. The horizontal dashed grey line marks zero correlation. The lower panels show the corresponding p-values from one-sample tests assessing whether the subject-level correlations were significantly greater than zero. P-values are displayed on a logarithmic scale, and the red dashed line indicates p = 0.05.

**Figure S8.**
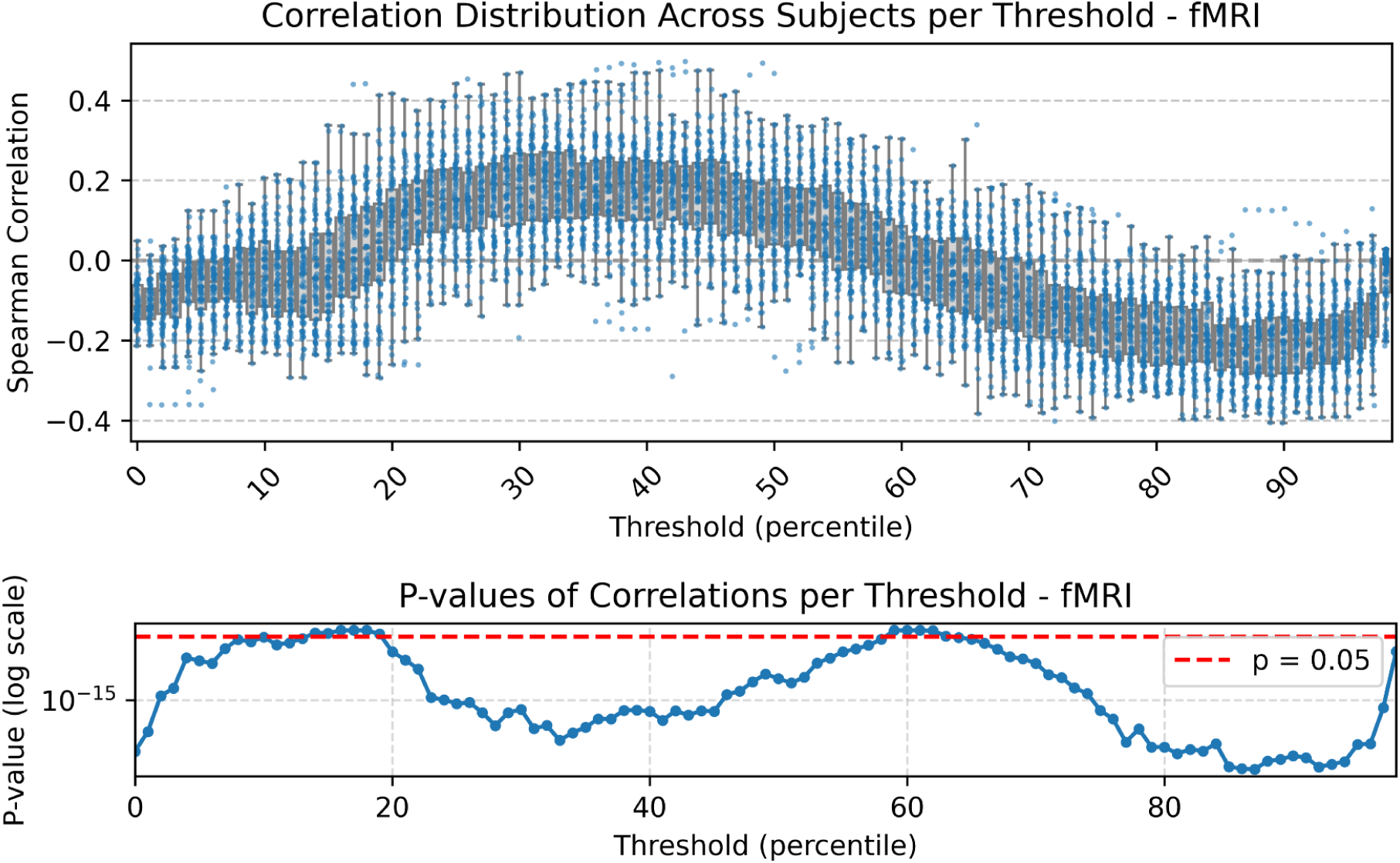
Structure-function correlation: validity across different thresholds, one-sample t-test at the population level (fMRI). Same analysis as in Figure S7 for MEG and EEG, but applied to the fMRI dataset. For each threshold, the Spearman correlation between regional flexibility-gradient values and regional structural connectivity strength was computed separately for each subject. The upper panel shows the distribution of subject-level correlations across thresholds, while the lower panel reports the corresponding p-values from one-sample tests against zero. P-values are shown on a logarithmic scale, and the red dashed line indicates p = 0.05.

**Figure S9:**
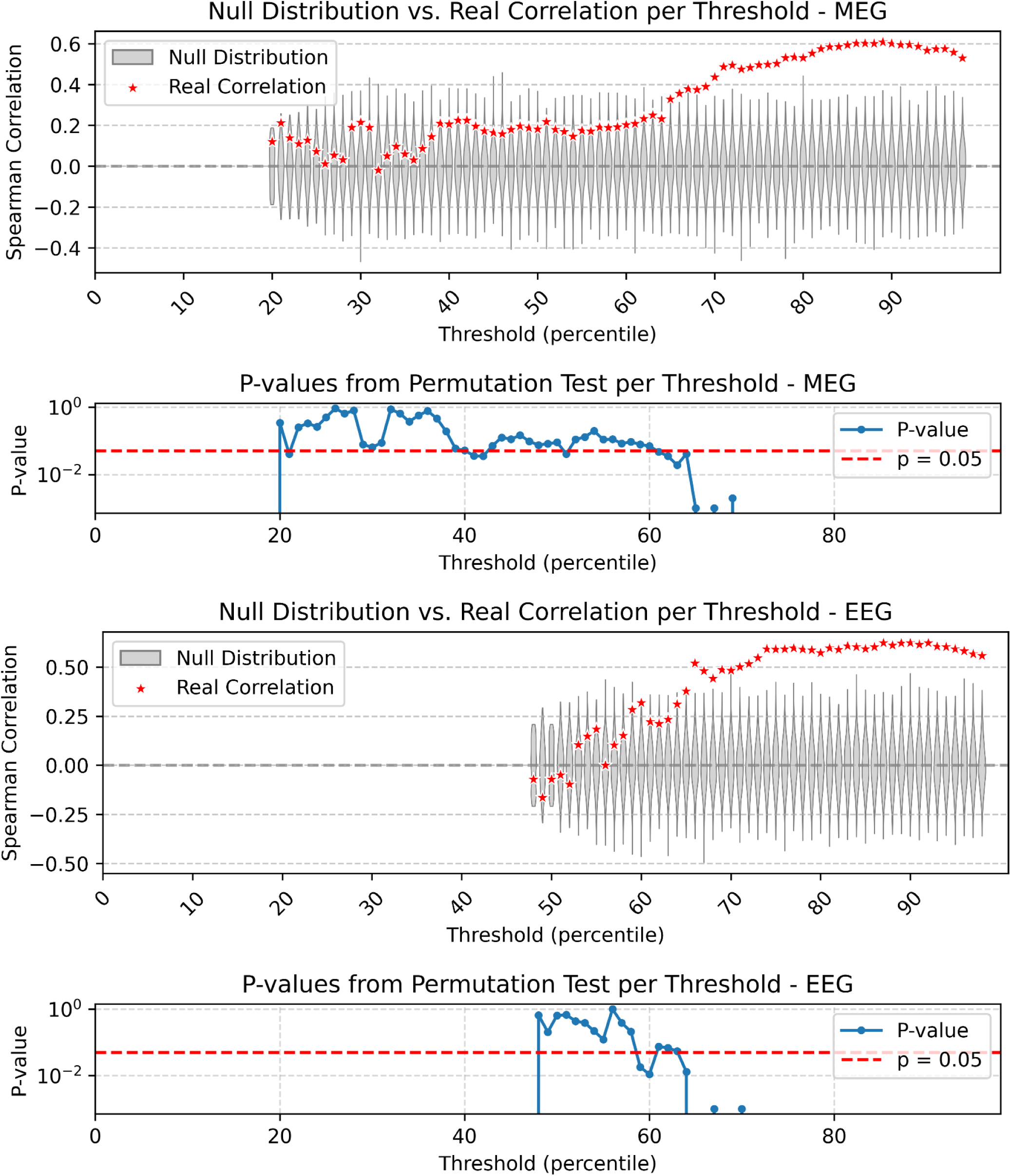
Structure-function correlation: validity across different thresholds, null model (EEG - MEG). This figure evaluates the robustness of the relationship between structural connectivity strength and the flexibility gradient (FG) in MEG and EEG data using a permutation null model (as in figure S4). For each threshold, the empirical Spearman correlation between regional FG values and regional structural strength is shown with red stars. The grey distributions represent null correlations obtained after cyclically permuting the regional data before recomputing the correlation, thereby disrupting the original correspondence between structural strength and FG while preserving the structure of the data. The lower panels show the corresponding permutation-test p-values across thresholds, with the red dashed line indicating p = 0.05.

**Figure S10:**
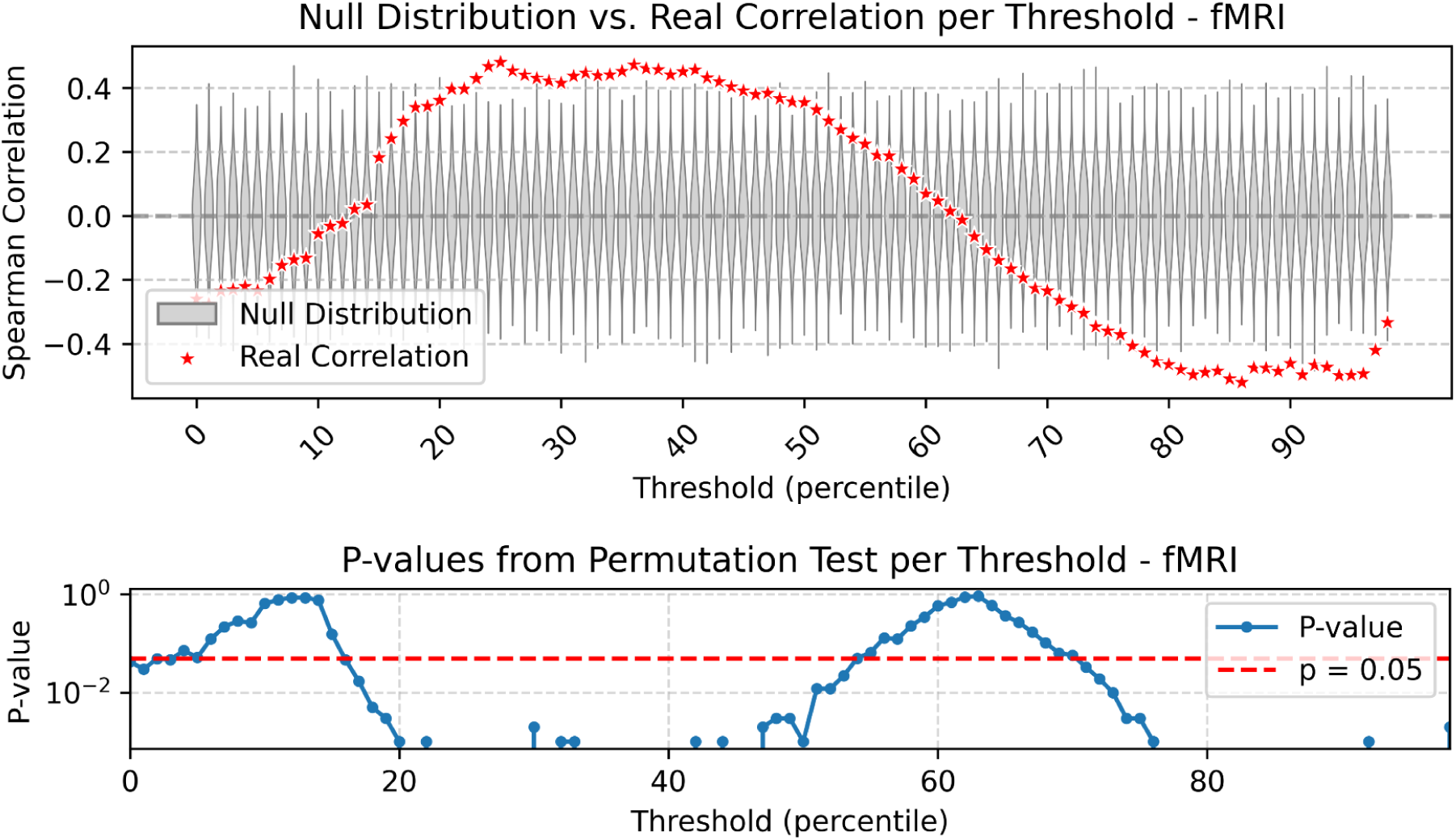
Structure-function correlation: validity across different thresholds, null model (fMRI). Same analysis as in Figure S9 for MEG and EEG, but applied to the fMRI dataset. For each threshold, the empirical Spearman correlation between regional flexibility-gradient values and regional structural connectivity strength is shown with red stars. The grey distributions represent null correlations obtained after randomly permuting the regional data before recomputing the correlation. The lower panel shows the corresponding permutation-test p-values across thresholds, displayed on a logarithmic scale. The red dashed line indicates p = 0.05.

**Figure S11:**
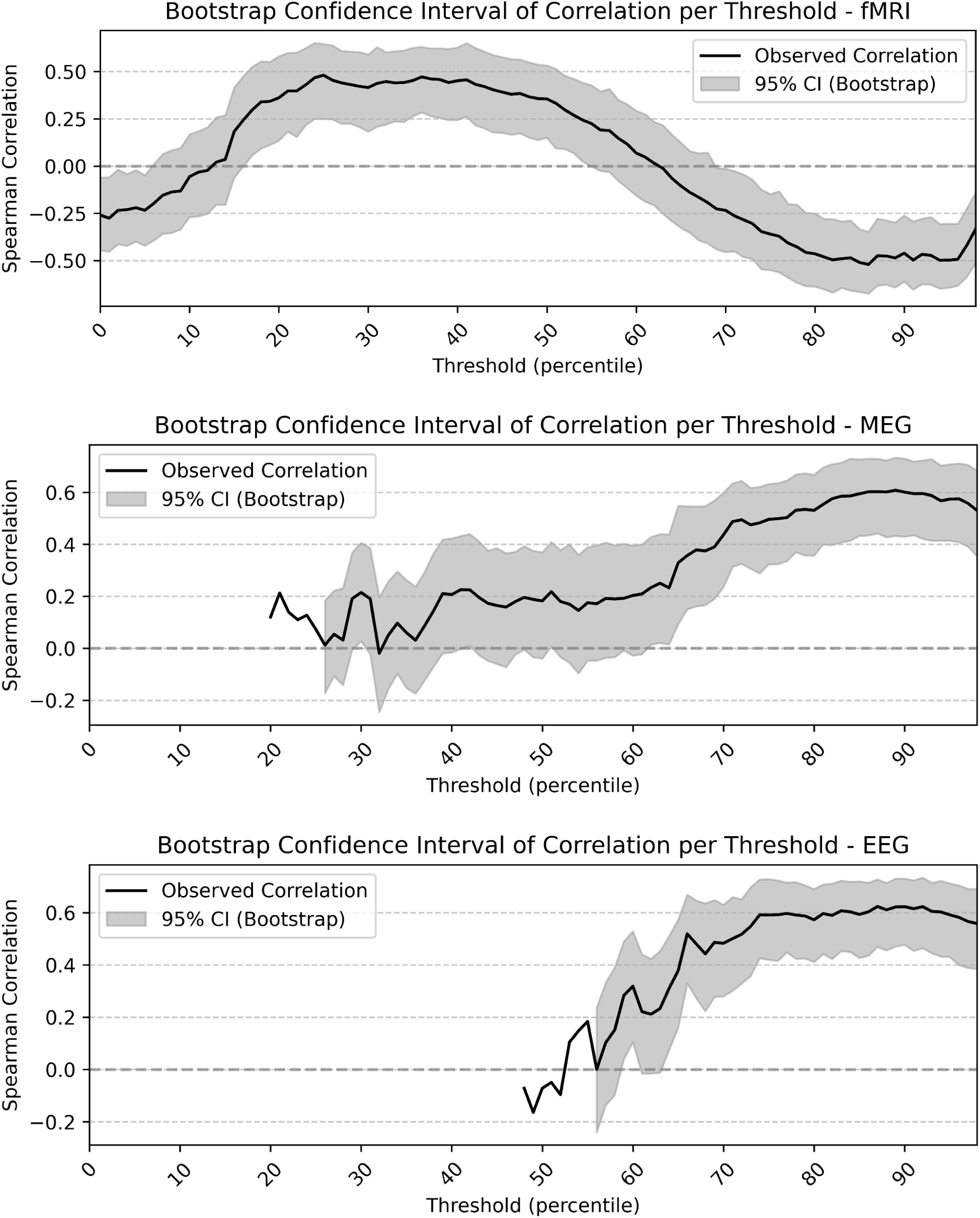
Structure-function correlation: validity across different thresholds, bootstrap. This figure shows the stability of the relationship between structural connectivity strength and the flexibility gradient (FG) across different binarisation thresholds for fMRI, MEG, and EEG data. For each threshold, the observed Spearman correlation between regional FG values and regional structural strength is shown as a black line. The grey shaded area represents the 95% confidence interval obtained by bootstrap resampling. This analysis assesses whether the observed structure–function correlations are robust to resampling of brain regions across thresholds.

## Additional analysis testing whether FG is explained by regional activation counts

Finally, we assessed whether the FG of brain regions is trivially driven by their number of events above threshold. To do so, we correlated FG for different thresholds with the sum across time of the binarized data. For EEG and MEG, this surrogate measure did not yield any significant statistical effects. In contrast, for fMRI we found that the overall activation energy trivially explained the FG at very low and very high thresholds. This likely reflects the short duration of the recordings, i.e., the limited number of time points, such that, at extreme thresholds, the FG becomes dominated by local activation counts (the number of samples exceeding threshold in each region). For this reason, we restricted our main analysis to thresholds between the 40th and 60th percentiles, where the correlation between number of activation and FG was not significant for one or more statistical tests.

**Figure S12:**
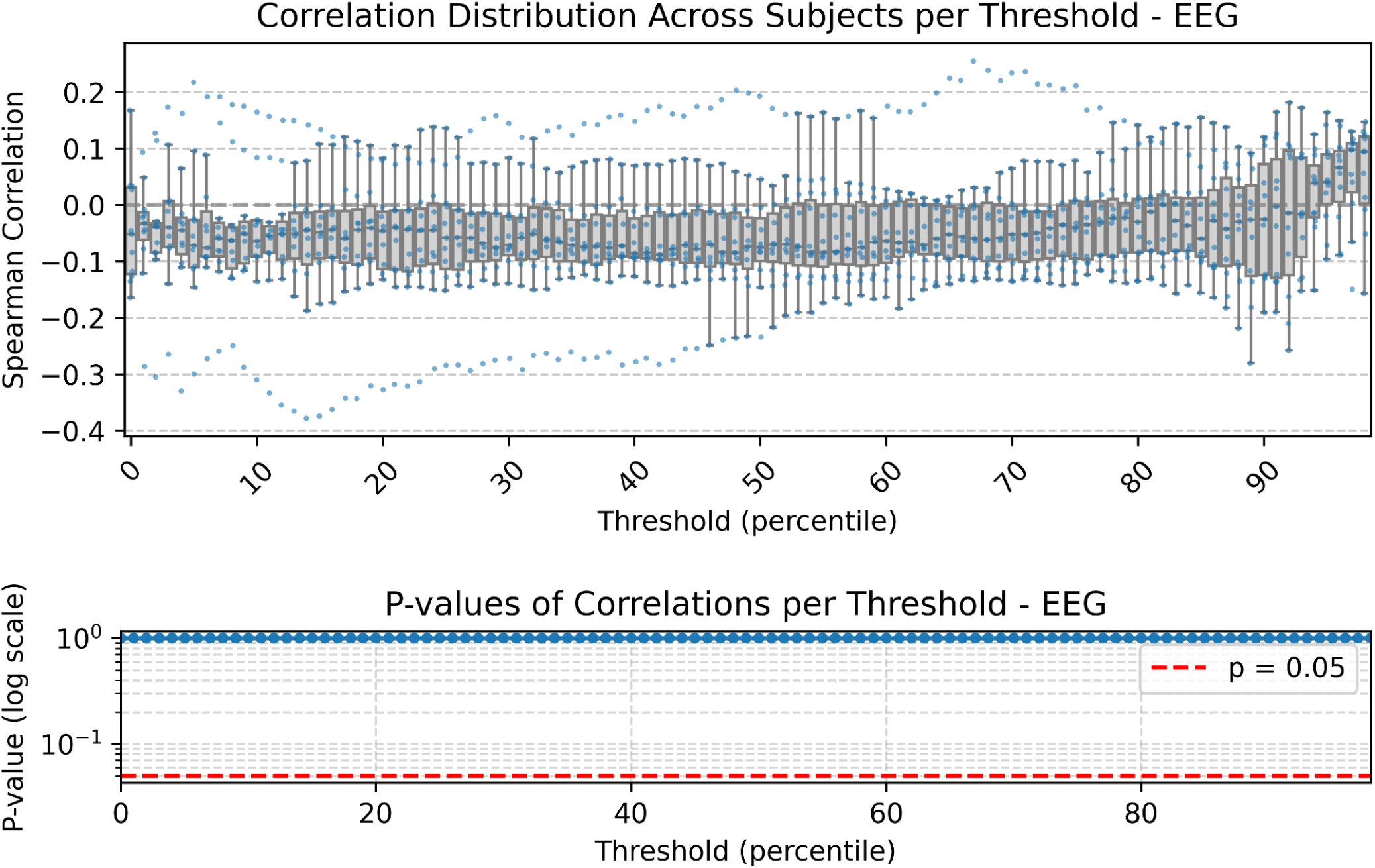
sums across time - FG correlation: validity across different thresholds, one-sample t-test at the population level (EEG). This figure tests whether the flexibility gradient (FG) is trivially explained by the number of supra-threshold activations in each brain region. For each threshold, we computed the Spearman correlation between regional FG values and the sum across time of the binarized EEG activity. The upper panel shows the distribution of subject-level correlations across thresholds, while the lower panel reports the corresponding p-values from one-sample tests against zero. P-values are shown on a logarithmic scale, and the red dashed line indicates p = 0.05. Across thresholds, activation counts do not show a consistent significant relationship with FG in EEG.

**Figure S13:**
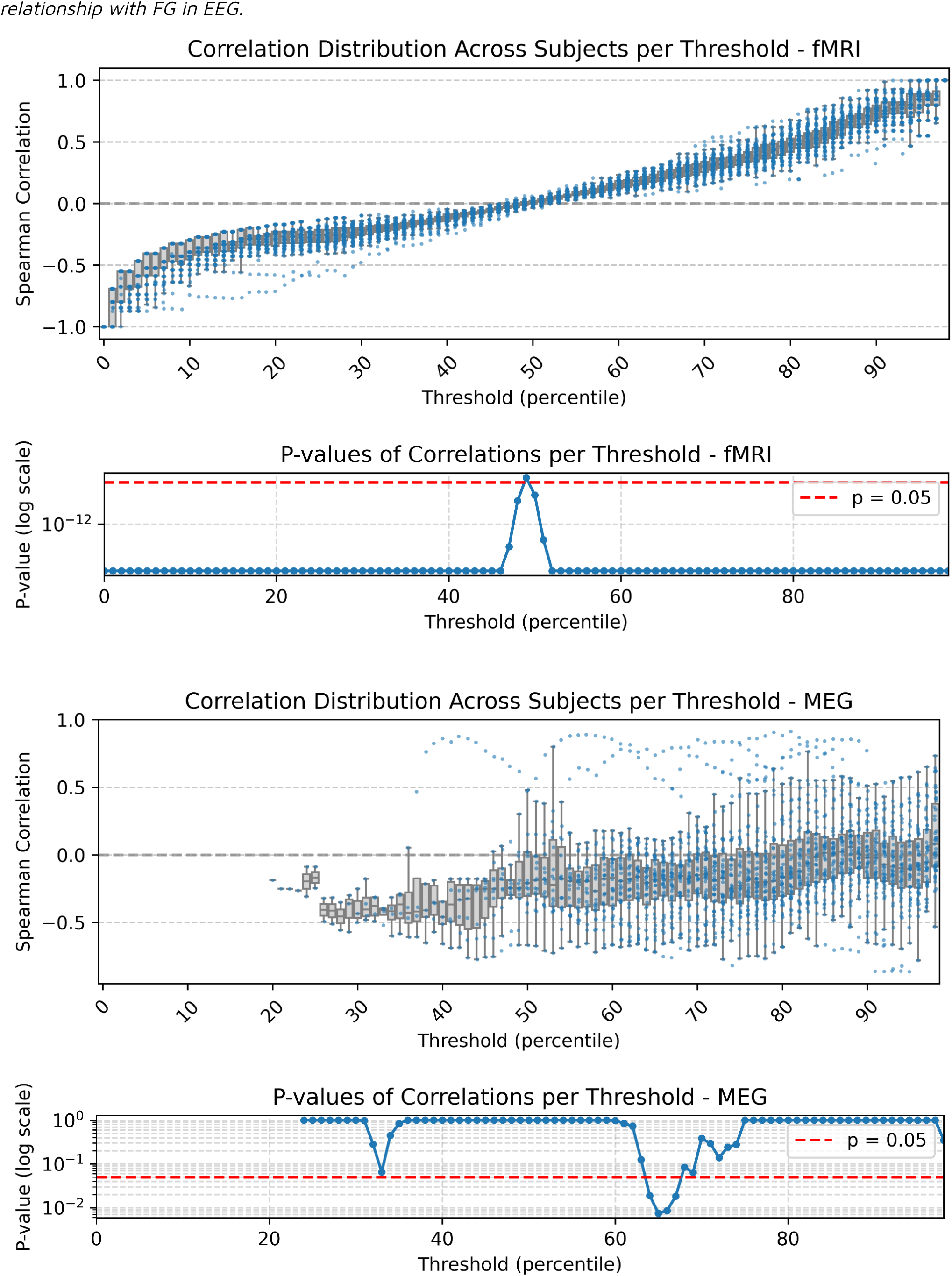
sums across time - FG correlation: validity across different thresholds, one-sample t-test at the population level (fMRI - MEG). Same analysis as in Figure S12, but applied to the fMRI and MEG datasets. For each threshold, the Spearman correlation between regional FG values and the number of supra-threshold activations was computed separately for each subject. The upper panels for each modality show the distribution of subject-level correlations across thresholds, and the lower panels show the corresponding p-values from one-sample tests against zero. While MEG does not show a consistent significant relationship between activation counts and FG, fMRI shows significant correlations at low and high thresholds, indicating that FG can be partially driven by regional activation counts at extreme fMRI thresholds.

**Figure S14:**
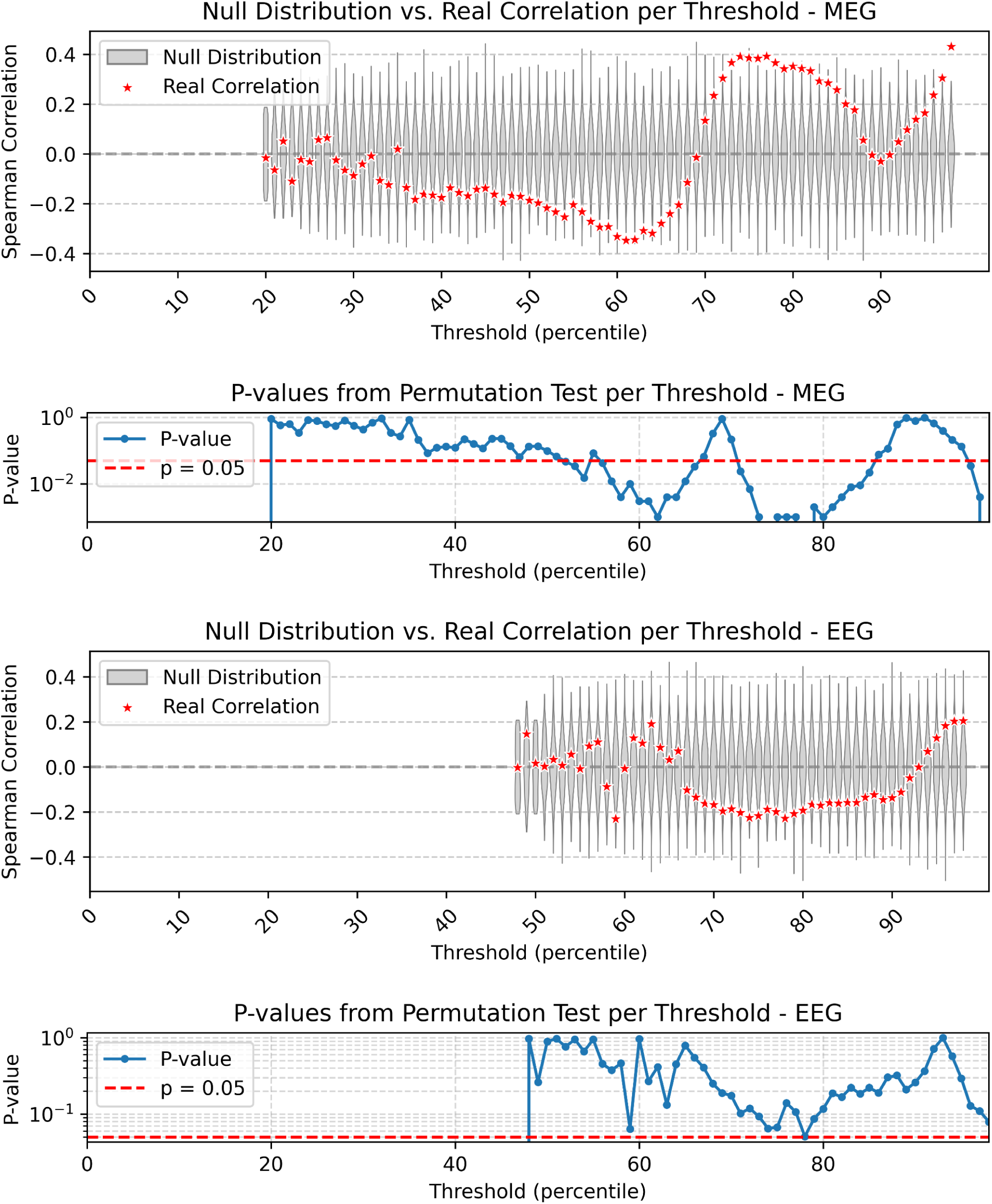
sums across time - FG correlation: validity across different thresholds, null model (MEG - EEG). This figure evaluates whether the relationship between activation counts and FG exceeds what would be expected under a cyclic-permutation null model. For each threshold, the empirical Spearman correlation between regional FG values and the number of supra-threshold activations is shown with red stars. The grey distributions represent null correlations obtained after cyclically permuting the regional data before recomputing the correlation. The lower panels show the corresponding permutation-test p-values across thresholds, with the red dashed line indicating p = 0.05. The results show that activation counts do not consistently explain FG in MEG or EEG.

**Figure S15:**
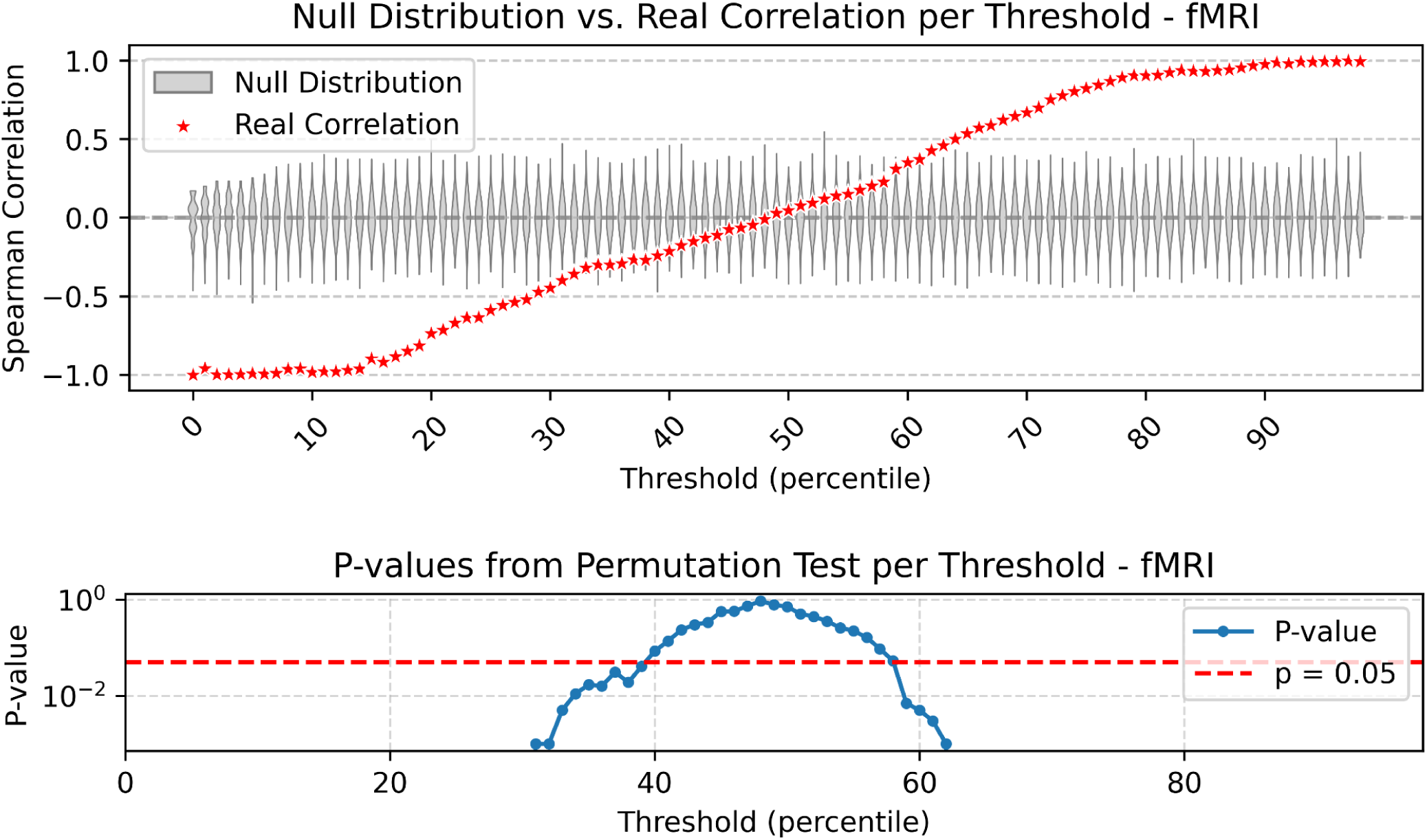
sums across time - FG correlation: validity across different thresholds, null model (fMRI). Same analysis as in Figure S14, but applied to the fMRI dataset. For each threshold, the empirical Spearman correlation between regional FG values and the number of supra-threshold activations is shown with red stars. The grey distributions represent null correlations obtained after cyclically permuting the regional data before recomputing the correlation. The lower panel shows the corresponding permutation-test p-values across thresholds, displayed on a logarithmic scale. The results indicate that activation counts significantly explain FG at very low and very high fMRI thresholds, supporting the restriction of the main fMRI analyses to intermediate thresholds.

**Figure S16:**
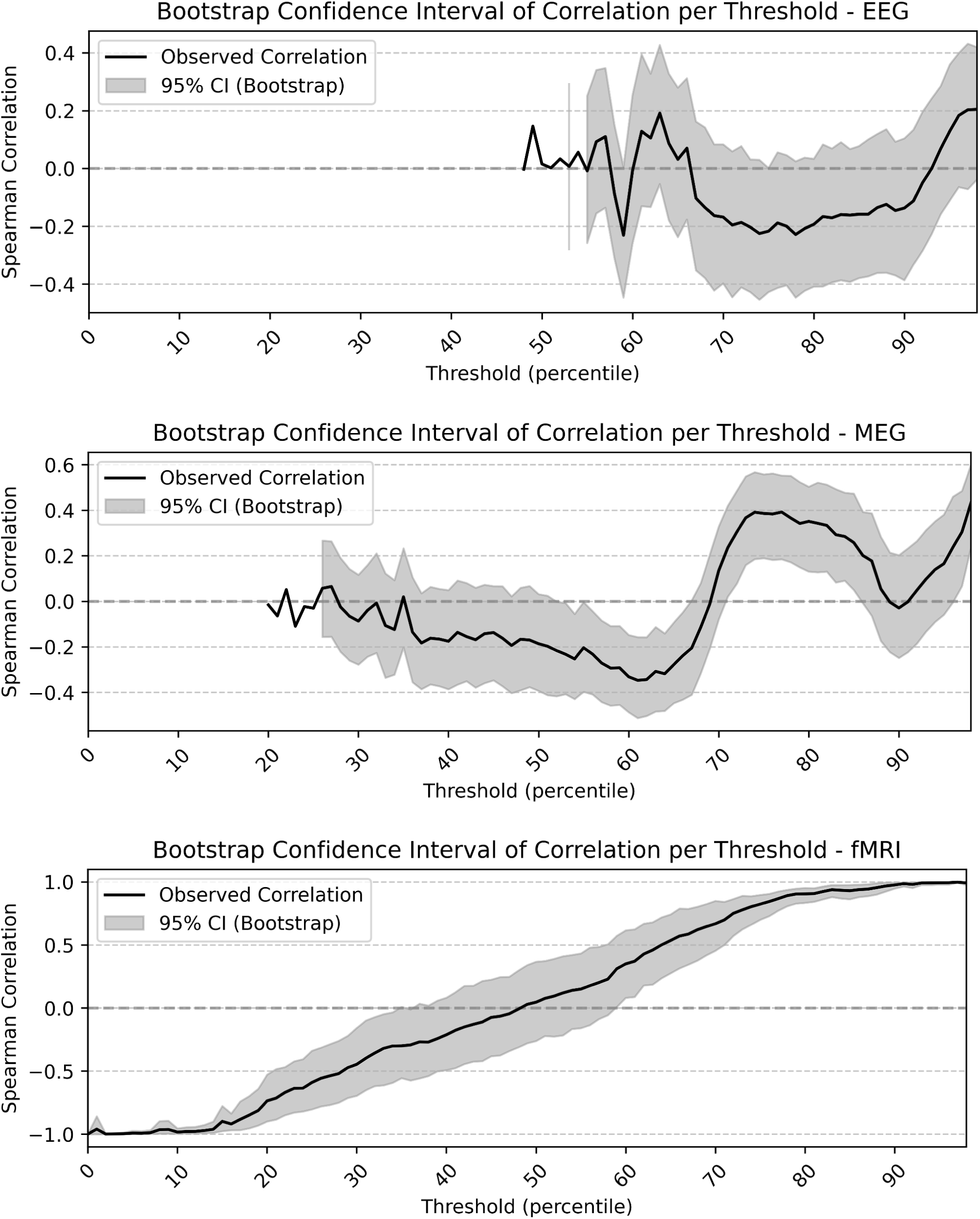
sums across time - FG correlation: validity across different thresholds, bootstrap. This figure shows the robustness of the relationship between activation counts and FG across thresholds for EEG, MEG, and fMRI. For each threshold, the observed Spearman correlation between regional FG values and the number of supra-threshold activations is shown as a black line. The grey shaded area represents the 95% confidence interval obtained by bootstrap resampling. For EEG and MEG, the relationship between activation counts and FG is not consistently positive or robust across thresholds. In contrast, fMRI shows a strong threshold-dependent pattern, with activation counts explaining FG at extreme thresholds but not within the intermediate range used for the main analyses.

## Notes

### Competing Interest Statement

The authors have declared no competing interest.

### Summary of Updates

In this revised version, we have reframed the main message of the manuscript, incorporated additional relevant references, and added new paragraphs to the Introduction and Discussion sections. We have also expanded the supplementary materials to include more detailed explanations.

